# NADPH and glutathione redox link TCA cycle activity to endoplasmic reticulum stress

**DOI:** 10.1101/822775

**Authors:** Erica R. Gansemer, Kyle S. McCommis, Michael Martino, Abdul Qaadir King-McAlpin, Matthew J. Potthoff, Brian N. Finck, Eric B. Taylor, D. Thomas Rutkowski

## Abstract

Endoplasmic reticulum (ER) stress is associated with dysregulated metabolism, but little is known about how the ER responds to metabolic activity. Here, working primarily in mouse hepatocytes, we show that decreasing the availability of substrate for the TCA cycle diminished NADPH production and attenuated ER stress in a manner that depended on glutathione oxidation. ER stress was also alleviated by impairing either TCA-dependent NADPH production or Glutathione Reductase. Conversely, stimulating TCA activity favored NADPH production, glutathione reduction, and ER stress. Validating these findings, we show that deletion of the mitochondrial pyruvate carrier, which is known to decrease TCA cycle activity and protect the liver from diet-induced injury, also diminished NADPH, elevated glutathione oxidation, and alleviated ER stress. These results provide independent genetic evidence that mitochondrial oxidative metabolism is linked to ER homeostasis. Our results demonstrate a novel pathway of communication between mitochondria and the ER, through relay of redox metabolites.

## Introduction

As the gateway to the secretory pathway, the ER must properly synthesize and fold nascent secretory and membrane proteins. Disruption of this process, known as ER stress, is associated with many disease states, including obesity and its comorbidities (Mohan, R, Brown, Ayyappan, & G, 2019). Thus, it is important to understand how ER homeostasis is perturbed, particularly by metabolic disruption. The ER and the mitochondrial network, although not connected to each other by secretory pathway traffic, are intertwined both physically and functionally. The ER makes close physical contacts with mitochondria to facilitate the exchange of metabolites (Raffaello, Mammucari, Gherardi, & Rizzuto, 2016; Vance, 2014; Yoboue, Sitia, & Simmen, 2018). The ER also communicates with mitochondria via signaling from the unfolded protein response (UPR), which is activated by ER stress and signals through the three ER-resident stress sensors IRE1, PERK, and ATF6 (Walter & Ron, 2011). The UPR regulates mitochondrial activity at several levels, including enhancing mitochondrial protein quality control, augmenting ER-mitochondrial interactions and calcium signaling, and contributing to mitochondrial depolarization and initiation of apoptosis (Y. Fan & Simmen, 2019; Gutierrez & Simmen, 2018; Rainbolt, Saunders, & Wiseman, 2014). While the pathways through which ER stress and the UPR regulate mitochondrial function are becoming clearer, the converse—how mitochondrial function impacts ER homeostasis—is less understood.

In the liver, the UPR suppresses a large number of genes involved in metabolic processes (Rutkowski et al., 2008). One of the processes suppressed is fatty acid β-oxidation (DeZwaan-McCabe et al., 2017), which takes place in mitochondria. The role of the UPR is to restore ER function during stress, suggesting that suppressing β-oxidation in the mitochondria is functionally beneficial to achieving ER homeostasis. However, it is not clear how metabolic activity within the mitochondria might be conveyed to the ER.

The impact of metabolic activity on ER homeostasis is most evident form the association of ER stress with obesity—particularly in highly metabolically active tissues such as liver, pancreas, and adipose (Cnop, Foufelle, & Velloso, 2012). Lipotoxicity (i.e., damage caused by the inappropriate accumulation of lipids in non-adipose tissue), inflammation, and oxidative stress have all been shown to contribute to obesity-associated ER stress (Fu, Watkins, & Hotamisligil, 2012; Salvado, Palomer, Barroso, & Vazquez-Carrera, 2015). Yet, independent of diet content, feeding after a fast is sufficient to elicit ER stress in the liver (Gomez & Rutkowski, 2016; Oyadomari, Harding, Zhang, Oyadomari, & Ron, 2008; Pfaffenbach et al., 2010) and to alter the extent of physical contacts between the ER and mitochondria (Theurey et al., 2016). These findings suggest that ER homeostasis is acutely and intrinsically connected with metabolism even apart from the problems brought on by obesity. However, the biochemical pathways linking catabolism to ER function are not known.

The tricarboxylic acid (TCA) cycle is the central hub of metabolism, participating in both catabolism and anabolism. Acetyl-CoA enters the cycle after either oxidative breakdown of lipids within the mitochondria or after conversion of pyruvate, generated from glycolysis in the cytosol, by the mitochondrial pyruvate dehydrogenase complex. Canonically, TCA activity yields NADH and FADH_2_ for the electron transport chain. However, the TCA cycle also provides precursors for biosynthetic pathways (Owen, Kalhan, & Hanson, 2002). In addition, the cycle can produce NADPH in addition to NADH, due to the activity of isozymes that reside either in the mitochondria (isocitrate dehydrogenase 2/IDH2) or cytosol (IDH1 or malic enzyme/ME1) (Rydstrom, 2006). Therefore, activity of the TCA cycle is likely to influence cellular processes by mechanisms beyond just the production of ATP from the electron transport chain.

NADPH is used as a cofactor by glutathione reductase (GR) to reduce oxidized glutathione (GSSG→2GSH), and likewise by thioredoxin reductase to reduce oxidized thioredoxin. Both of these molecules contribute to defense against oxidative stress (Sies, Berndt, & Jones, 2017), and both have connections to ER protein biogenesis. Thioredoxin has been shown to be a source of electrons for reduction and isomerization of disulfide bonds of ER client proteins (Poet et al., 2017). The oxidized form of glutathione (GSSG), which predominates in the ER lumen compared to the cytosol, was formerly thought to reoxidize protein disulfide isomerase (PDI) to promote ER disulfide bond formation. However, since the discovery of alternative pathways for PDI reoxidation (Frand & Kaiser, 1999; Tu, Ho-Schleyer, Travers, & Weissman, 2000; Zito, Melo, et al., 2010), the role of glutathione in the ER is now much less clear (Delaunay-Moisan, Ponsero, & Toledano, 2017; Tsunoda et al., 2014). Whether elevated GSSG might be beneficial to ER function under some cellular conditions but detrimental in others is also unknown.

Despite the centrality of the TCA cycle to cellular function and hints that its activity might be tied to ER stress (Mogilenko et al., 2019; Xin et al., 2018), its involvement in ER homeostasis has not been investigated. Here, we used primary hepatocytes and other metabolically active cell types to investigate the relationship between metabolic activity and ER stress. We show that TCA cycle activity links lipid and carbohydrate catabolism to ER homeostasis through production of NADPH and redox regulation of glutathione. Our findings delineate a novel mechanism of communication from mitochondria to the ER, reveal an unexpected protective role for GSSG, and provide a plausible pathway by which ER homeostasis is linked to metabolic activity.

## Results

### Inhibition of β-oxidation alleviates ER stress in metabolically active cell types

We have previously shown *in vivo* and in hepatoma cells *in vitro* that inhibiting β-oxidation either pharmacologically or genetically diminishes ER stress signaling (Tyra, Spitz, & Rutkowski, 2012). In order to identify the pathway by which β-oxidation and ER homeostasis are linked, we first asked whether etomoxir, which blocks β-oxidation by inhibiting the CPT1-dependent transport of fatty acyl-CoAs into the mitochondria for oxidation (Weis, Cowan, Brown, Foster, & McGarry, 1994; Yao et al., 2018), could diminish ER stress signaling in primary hepatocytes *in vitro* as it does in the liver *in vivo*. Treatment of primary hepatocytes with the ER stressor tunicamycin (TM) upregulated UPR-responsive mRNAs, whereas etomoxir cotreatment suppressed this effect (Figure 1A). Dampened ER stress signaling was also evident from diminishment of the splicing of the IRE1α nuclease target *Xbp1* (Figure 1B) and of the upregulation of the stress-regulated factor CHOP (Figure 1C). Because these events are differentially regulated by the three limbs of the UPR, our data suggest that etomoxir diminishes signaling from all three UPR pathways. This result is consistent with our previous findings, in which ER stress signaling in hepatoma cells was diminished by etomoxir, by inhibition of β-oxidation with a separate agent, and by knockdown of CPT1 (Tyra et al., 2012).

**Figure 1:**
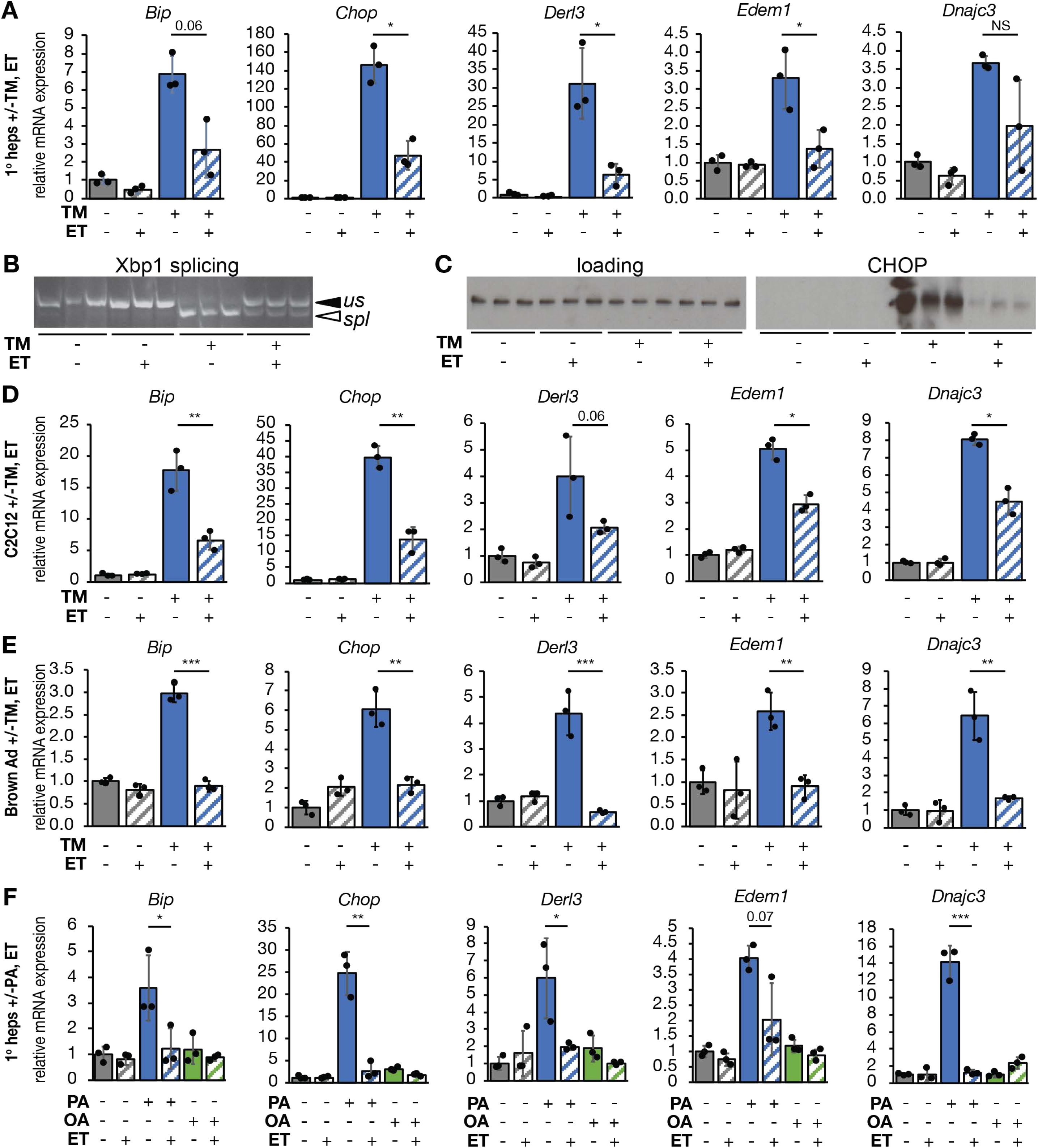
Inhibiting β-oxidation attenuates UPR activity in metabolically active cells and protects against different ER stressors. **(A-C)** Primary hepatocytes were treated with vehicle or 250 ng/mL tunicamycin (TM) in the presence or absence of 25 µg/mL etomoxir (ET) for 8 h. Activation of the UPR was assessed by qRT-PCR of a sampling of UPR target genes (A), splicing of *Xbp1* by conventional RT-PCR (B), and CHOP protein expression by immunoblot (C). In (B) and (C), and elsewhere, each lane represents an independently treated well. Loading control for immunoblot was calnexin, which does not change in response to TM or ET. **(D, E)** Same as except using C2C12 myoblasts (D) or primary brown adipocytes (E). **(F)** mRNA expression of UPR markers was assessed by qRT-PCR in primary hepatocytes treated with 200 µM palmitate (PA) or oleate (OA) and 25 µg/mL ET for 8 h. For this and all subsequent figures, error bars represent means ± S.D.M. *, p<0.05, **, p<0.01, ***, p<0.001 by two-tailed t-test and Benjamini-Hochberg adjustment for multiple comparisons. p-values between 0.05 and 0.1 are given as values. NS, p>0.1.

We next determined if the apparent protective effects of etomoxir were limited to one cell type or to one ER stressor. Using the selection of UPR-regulated mRNAs in Figure 1A as a broad indicator of ER stress signaling, we found that etomoxir also diminished UPR activation in C2C12 myoblasts (Figure 1D), and immortalized (data not shown) and primary (Figure 1E) brown adipocytes. Hepatocytes, myocytes, and brown adipocytes are characterized by particularly high lipid metabolic activity (Frayn, Arner, & Yki-Jarvinen, 2006). In contrast, etomoxir had no significant effect on the expression of UPR target genes in primary mouse embryonic fibroblasts (Figure S1). In primary hepatocytes subjected to ER stress by palmitate loading, etomoxir likewise diminished ER stress signaling (Figure 1G). (As expected, the unsaturated fatty acid oleate did not cause ER stress.) Palmitate is thought to elicit ER not by disrupting protein folding *per se* but by altering ER membrane fluidity, activating the UPR stress sensors through their transmembrane domains (Volmer, van der Ploeg, & Ron, 2013). Thus, etomoxir diminishes ER stress induced by stressors that act by divergent mechanisms.

The effects of etomoxir on ER stress signaling could be due to improvement of ER homeostasis in some way, or to simple inhibition of UPR signaling. We did not observe any robust differences in the efficiency with which ER client proteins were secreted in hepatocytes treated with etomoxir compared to those that were not treated (data not shown). However, as a separate assessment of ER homeostasis, we examined ER ultrastructure in hepatocytes treated with TM, with or without etomoxir. Consistent with previous reports (Finnie, 2001; Rutkowski et al., 2006), TM elicited marked ER dilation accompanied by a loss of lamellar structure and overall disorganization, although some areas of grossly normal ER were present and were predominantly juxtaposed near mitochondria. In contrast, these disruptions were largely prevented by cotreatment with etomoxir (Figure 2A) as confirmed by two independent, blinded scorers (Figure 2B). Although etomoxir diminished ER stress signaling, it did not block it entirely. Higher doses of TM elicited UPR activation in etomoxir-treated cells to an extent comparable to that observed with lower doses in non-etomoxir-treated cells (Figure 2C). Therefore, the UPR in etomoxir-treated cells remained competent for signaling. Together, these results suggest that inhibiting β-oxidation alleviates ER stress.

**Figure 2:**
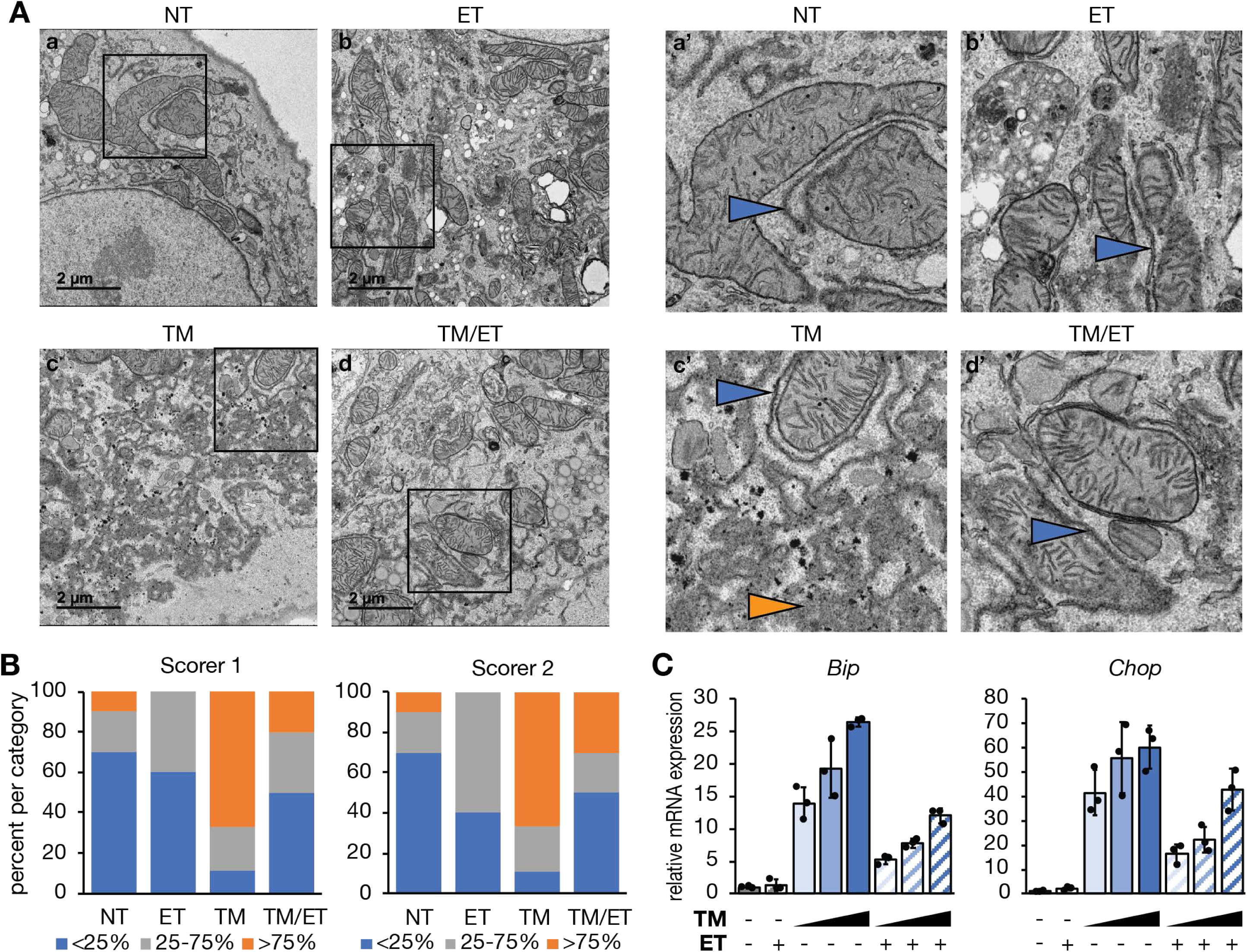
Inhibiting β-oxidation improves ER ultrastructure. **(A)** Primary hepatocytes were treated with vehicle, 250 ng/mL TM, 25 µg/mL ET, or TM and ET for 8 h and fixed in 2.5% glutaraldehyde. Fixed cells were imaged by transmission electron microscopy. Inset panels are indicated. Blue arrowheads represent areas of structurally normal ER, while orange arrowhead represent dysmorphic ER. **(B)** Images were scored blindly and binned based on the percentage of “dilated” ER in the image. Normal (blue): <25%, Moderate (gray): 25-75%, and Severe (orange): >75%. n = 9-10 cell images per group. **(C)** Primary hepatocytes were treated with 25 μg/ml ET in the presence of 0.5, 2, or 10 μg/ml TM for 8 h, and expression of *Bip* and *Chop* was detected by qRT-PCR.

To provide a direct test of whether etomoxir could alleviate ER stress caused by the accumulation of misfolded protein, we examined its ability to reverse ER stress associated with overexpression of the null Hong Kong (NHK) protein. NHK is encoded by a nonsense mutation of α1-antitrpysin that results in a truncated product which is retained in the ER and undergoes ER-associated degradation, resulting in ER stress in diverse cell types (Nagasawa, Higashi, Hosokawa, Kaufman, & Nagata, 2007; Ordonez et al., 2013; Sifers, Brashears-Macatee, Kidd, Muensch, & Woo, 1988; J. Wu et al., 2007). We transduced primary hepatocytes with a recombinant adenovirus expressing NHK under doxycycline (Dox)-inducible control (Ad-TetOn-NHK). As a control, cells were transduced with adenovirus constitutively overexpressing GFP instead (Ad-GFP). No ER stress was observed in Ad-TetOn-NHK-transduced cells in the absence of Dox (not shown). As expected, Dox treatment elicited ER stress in cells transduced with Ad-TetOn-NHK, but not in cells expressing Ad-GFP. The stress induced by NHK was markedly diminished by etomoxir (Figure 3A). Overexpression of NHK disrupted ER structure, resulting in the appearance of structurally amorphous ER puncta that were diminished by etomoxir treatment (Figure 3B). This finding was again substantiated by blinded analysis of EM images (Figure 3C). In principle, etomoxir treatment could alleviate NHK-dependent ER stress by reducing protein synthesis. However, we found no evidence that global translation rates were altered by etomoxir (Figure S2). Consistent with this observation, we found that, 4 hours after NHK induction, steady-state expression of NHK did not differ in cells treated or not with etomoxir (Figure 3D). However, at later time points, we observed diminished steady state expression of NHK in etomoxir-treated cells, and this distinction was lost when degradation of NHK was blocked by the proteasomal inhibitor MG-132 (Figure 3D). While the mechanisms by which etomoxir enhances ERAD remain under investigation, these results provide a separate line of evidence that inhibiting β-oxidation improves ER homeostasis, perhaps by affecting misfolded protein clearance.

**Figure 3:**
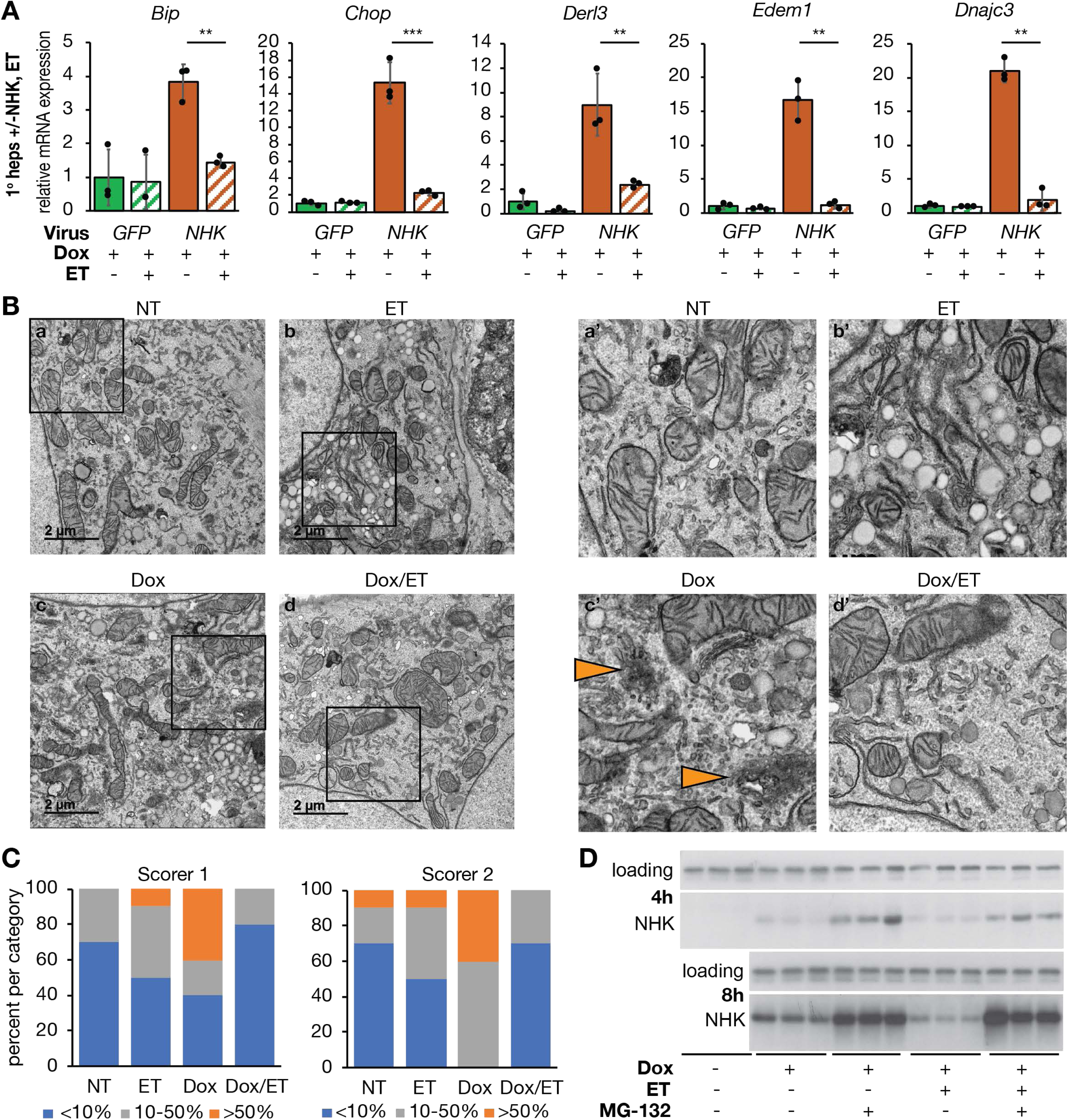
Inhibiting β-oxidation attenuates ER stress induced by overexpression of a constitutively misfolded protein. **(A)** Primary hepatocytes were infected with Ad-CMV-eGFP (control) or Ad-TetOn-NHK at an MOI of 1:1. NHK expression was induced with 500 ng/mL doxycycline (Dox) in the presence or absence of 25 µg/mL ET. mRNA expression of UPR markers was measured by qRT-PCR**. (B)** Primary hepatocytes infected with Ad-TetOn-NHK and treated with Dox and/or ET for 8 h were fixed in 2.5% glutaraldehyde for TEM. Orange arrowheads in insets denote dysmorphic ER. **(C)** Images were scored as described in Figure 2; n = 10 cell images per group. **(D)** Primary hepatocytes infected with Ad-TetOn-NHK were treated with Dox and ET as in (A) for 4 or 8 h in the presence or absence of 5 µM MG-132. NHK expression was analyzed by immunoblot. Loading control was calnexin.

### Inhibiting β-oxidation protects ER function through glutathione redox

We and others have previously shown that etomoxir raises the cellular ratio of oxidized (GSSG) to reduced (GSH) glutathione (Merrill et al., 2002; Pike, Smift, Croteau, Ferrick, & Wu, 2010; Tyra et al., 2012). Given the association of GSSG with the oxidative protein folding environment in the ER, we examined whether etomoxir could make the ER more oxidizing to proteins in the lumen. This was done by monitoring the oxidation of albumin, an endogenous hepatocyte ER client protein that forms 17 disulfide bonds (Peters & Davidson, 1982). We tested the resistance of albumin to reduction by treating cells with DTT, lysing them, and then modifying free sulfhydryls with PEG-maleimide, which retards migration upon SDS-PAGE (Winther & Thorpe, 2014). We found that treatment with DTT led to almost complete reduction of albumin in otherwise untreated cells, whereas etomoxir caused a significant increase in the preponderance of albumin that remained oxidized (Figure 4A). These results suggest that glutathione redox might mediate the protective effects of etomoxir.

**Figure 4:**
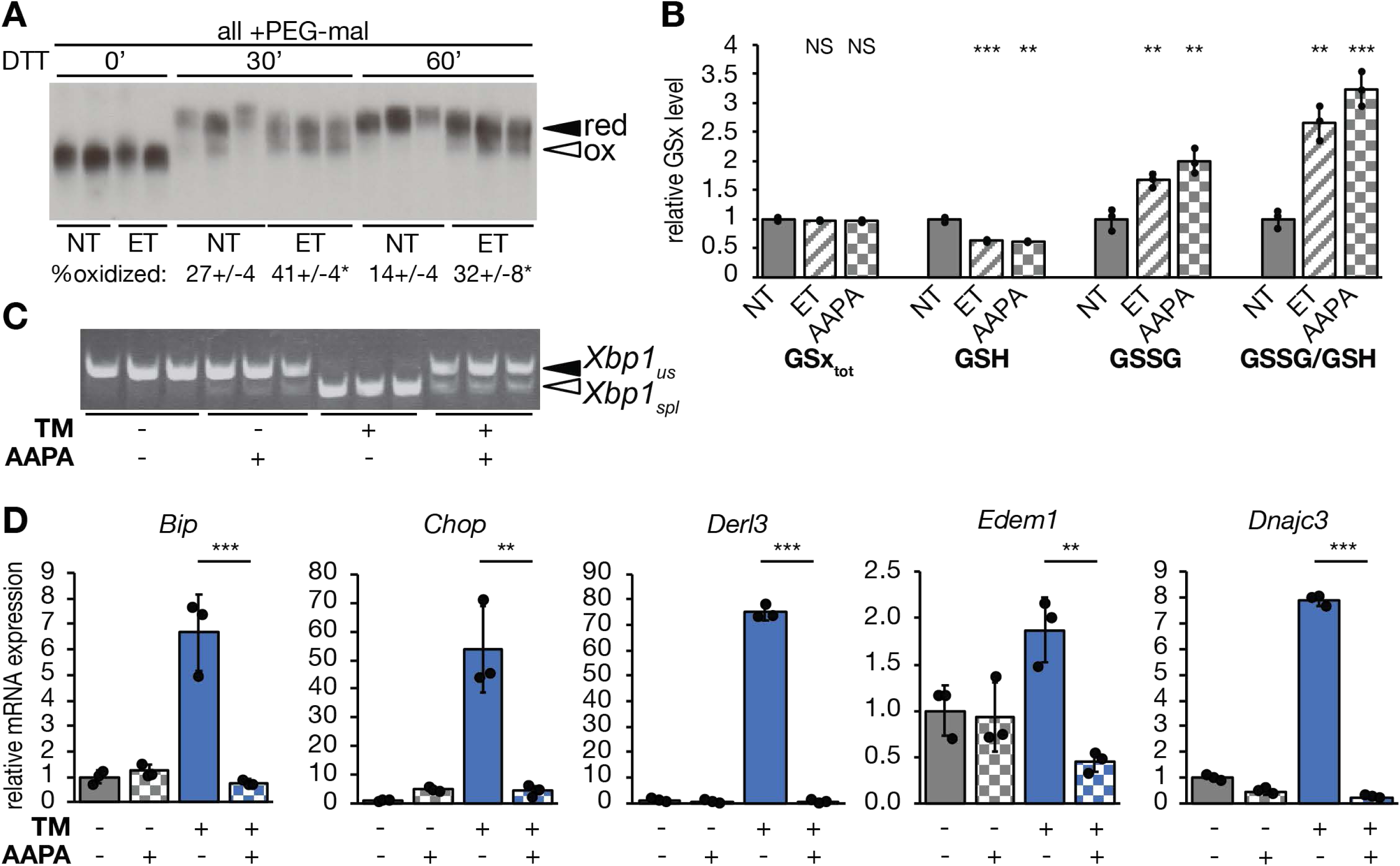
Inhibiting Glutathione Reductase attenuates ER stress. **(A)** Primary hepatocytes were treated with 25 µg/mL ET for 4 h followed by addition of 10 mM DTT for 0, 30, or 60 min. Protein lysates were treated with 4 µM mm(PEG)_24_ prior to SDS-PAGE and immunoblotting to detect endogenous albumin. The percentage of oxidized albumin is given below each group, with statistical comparison between ET-treated and non-treated cells. **(B)** Primary hepatocytes were treated with 25 μg/ml ET or 25 µM 2-AAPA for 8 h. Levels of total (GSx_tot_), reduced (GSH), and oxidized (GSSG) glutathione, and the ratio of GSSG to GSH were measured fluorometrically and expressed relative to untreated cells. **(C)** Splicing of *Xbp1* mRNA was measured by conventional RT-PCR, and **(D)** mRNA expression of UPR markers was measured by qRT-PCR in cells treated for 8 h with TM and/or 2-AAPA.

To first determine if glutathione redox plays a role in the protective effects of etomoxir, we used 2-AAPA to inhibit glutathione reductase (Seefeldt et al., 2009; Zhao et al., 2009), an enzyme with both mitochondrial and cytosolic activity that couples NADPH oxidation to GSSG reduction. We predicted that this treatment would phenocopy the protective effects of etomoxir. As expected, treatment with 2-AAPA diminished cellular GSH levels and increased GSSG, thus significantly elevating the GSSG/GSH ratio, similar to the effects of etomoxir (Figure 4B). Also similarly to etomoxir, 2-AAPA treatment alleviated ER stress, as determined by attenuated *Xbp1* splicing (Figure 4C) and diminished upregulation of UPR target genes (Figure 4D). Therefore, inhibiting glutathione reduction phenocopies the protective effects of etomoxir.

We next tested the converse prediction: that promoting glutathione reduction would cause ER stress. We treated hepatocytes with auranofin, which inhibits redox-active selenoproteins, including thioredoxin reductases and, at the dose used here, glutathione peroxidases (Chaudiere & Tappel, 1984; Roberts & Shaw, 1998; Scarbrough et al., 2012). In the liver, glutathione peroxidase 1 is two orders of magnitude more abundant than any thioredoxin reductase (Lai, Kolippakkam, & Beretta, 2008). Therefore, presumably due to its effects on glutathione peroxidase, auranofin treatment elevated cellular GSH levels and diminished GSSG, leading to a decrease in the GSSG/GSH ratio (Figure 5A). The relative resistance of albumin to reduction by DTT upon treatment with auranofin suggested that the ER environment was more reducing, whereas treatment with 2-AAPA made it more oxidizing (Figure 5B). Auranofin treatment alone was sufficient to induce a modest level of ER stress, seen in upregulation of UPR target genes in the absence of any other ER stress stimulus (Figure 5C). More importantly, auranofin completely blocked the ability of etomoxir to alleviate ER stress (Figure 5D). This epistatic relationship suggests that the protective effect of etomoxir requires glutathione oxidation.

**Figure 5:**
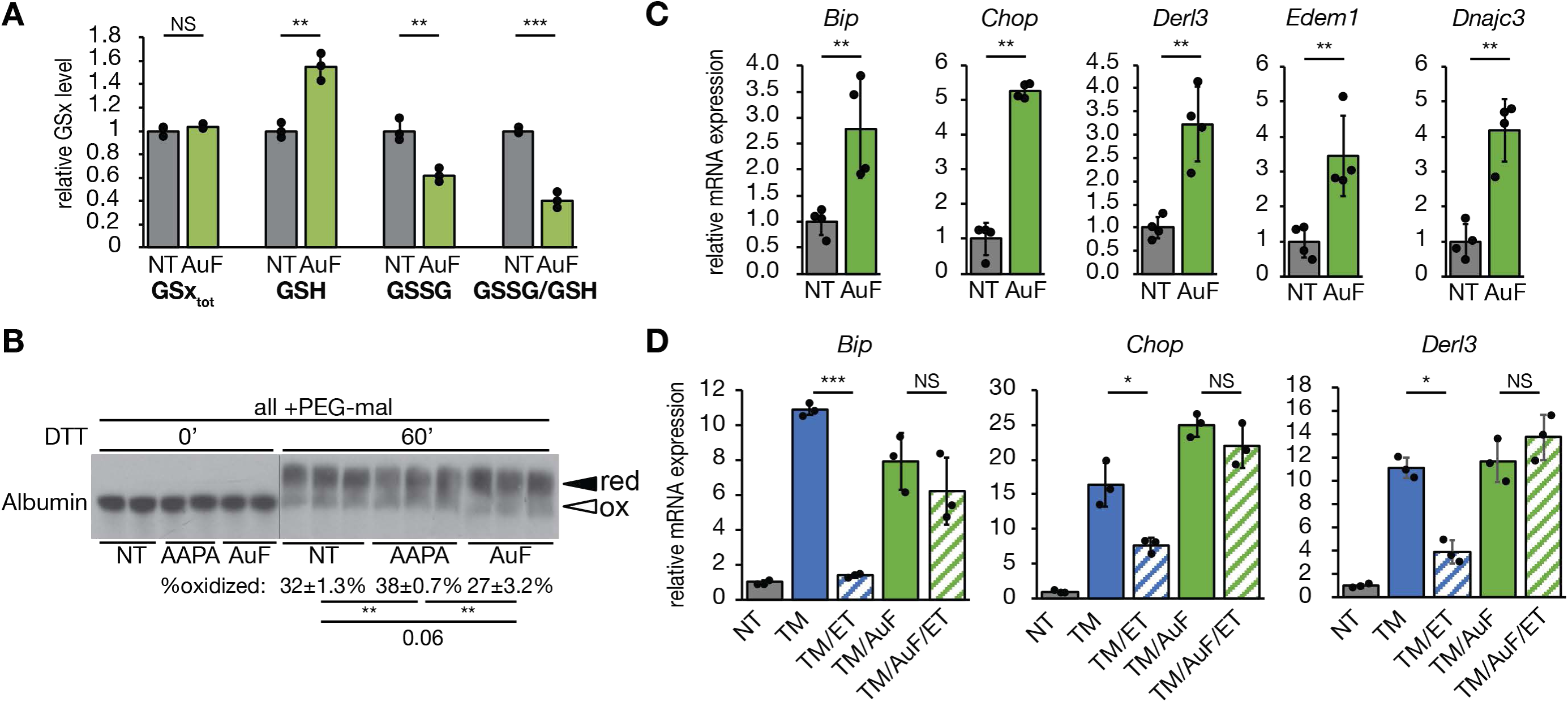
Glutathione oxidation is required for inhibition of β-oxidation to alleviate ER stress. **(A)** Levels of total, reduced, and oxidized glutathione were measured in primary hepatocytes treated with 10 µM auranofin (AuF) for 8 h. Glutathione was quantified as in Figure 4. **(B**) Hepatocytes were treated with 2-AAPA or AuF for 4 h prior to DTT and mm(PEG)_24_ as in Figure 4. **(C)** mRNA expression of UPR markers after treatment with AuF for 8h was quantified. **(D)** Hepatocytes were treated with TM, ET, and AuF as indicated. mRNA expression of UPR markers was measured by qRT-PCR.

### TCA cycle activity links oxidative metabolism to ER homeostasis

Although glutathione reductase uses NADPH to reduce glutathione, β-oxidation yields NADH and FADH_2_ but not NADPH. However, β-oxidation yields acetyl-CoA, which enters the TCA cycle by condensation with oxaloacetate for further oxidation. We therefore speculated that etomoxir might diminish production of NADPH from the TCA cycle, thereby resulting in elevated GSSG. In this model, it is not β-oxidation *per se* that is linked to ER homeostasis, but rather TCA cycle activity and the resultant production of NADPH. This model first predicted that etomoxir would diminish levels of both acetyl-CoA and NADPH—predictions which we then confirmed (Figure 6A, B). The model also predicted that impairing TCA-dependent NADPH production would elevate the GSSG/GSH ratio and alleviate ER stress, similarly to etomoxir treatment. Three TCA cycle isozymes generate NADPH: mitochondrial isocitrate dehydrogenase 2 (IDH2) and cytosolic IDH1 and malic enzyme (ME1). (The NADPH-producing mitochondrial ME3 is not expressed in the mouse liver). Using siRNAs, we specifically knocked down mRNA expression of each of these (Figure 6C). Knockdown of each gene diminished GSH levels, and knockdown of *Idh2* and *Me1* also increased GSSG levels, thus elevating the GSSG/GSH ratio (Figure 6D). Further, ER stress was diminished by knockdown of each of the three genes (Figure 6E).

**Figure 6:**
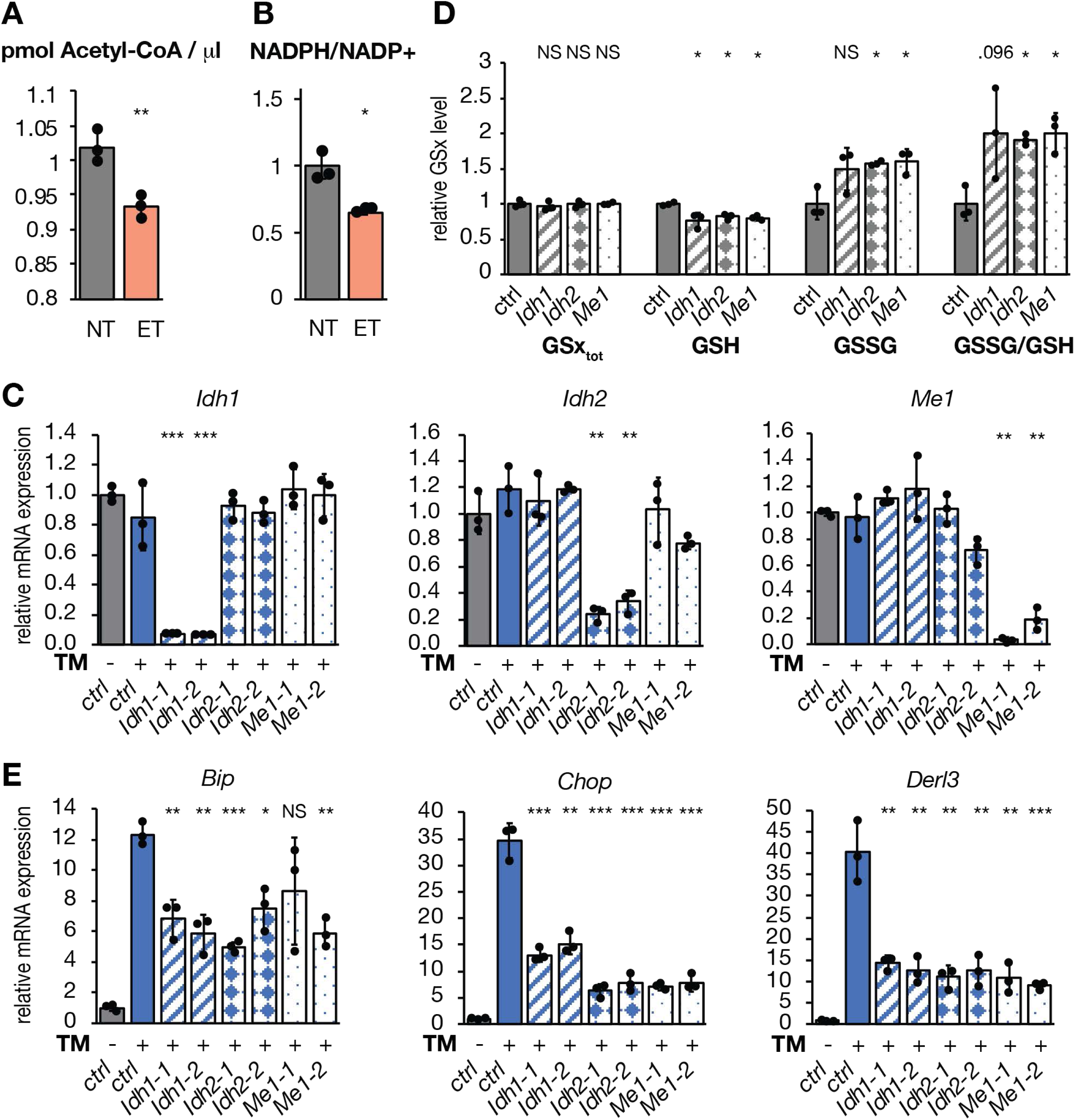
Diminishing TCA-dependent production of NADPH alleviates ER stress. **(A)** Acetyl-CoA levels or **(B)** the NADPH/NADP+ ratio were measured in primary hepatocytes treated with ET for 8 h and is expressed relative to untreated cells. **(C)** *Idh1*, *Idh2*, and *Me1* were knocked down with dsiRNA prior to treatment with TM for 8 h. A non-targeting dsiRNA was used as a control. Relative mRNA expression of *Idh1*, *Idh2*, and *Me1* was measured by qRT-PCR. **(D)** Glutathione was quantified in cells with knocked down *Idh1*, *Idh2*, or *Me1* (first oligo pair from Figure 6C in each case). **(E)** Expression of UPR target genes was assessed in cells from (C, D).

The Pyruvate Dehydrogenase complex oxidizes pyruvate produced by glycolysis to acetyl-CoA, and is a major control point of TCA cycle activity for carbohydrate metabolism (Gray, Tompkins, & Taylor, 2014). Pyruvate Dehydrogenase Kinases (PDKs) phosphorylate and inactivate this enzyme complex. To determine the effects of stimulating TCA cycle activity, we treated hepatocytes with dichloroacetate, which stimulates substrate entry into the TCA cycle by inhibiting PDKs (Constantin-Teodosiu, Simpson, & Greenhaff, 1999; C. Y. Wu et al., 2018). Dichloroacetate treatment substantially elevated the NADPH/NADP+ ratio (Figure 7A). In addition, dichloroaceetate treatment alone caused ER stress to an extent that was nearly as robust as the bona fide ER stressor TM, as seen by upregulation of UPR target genes (Figure 7B) and splicing of *Xbp1* (Figure 7C). Therefore, these data provide direct evidence that mitochondrial oxidative catabolic activity causes ER stress.

**Figure 7:**
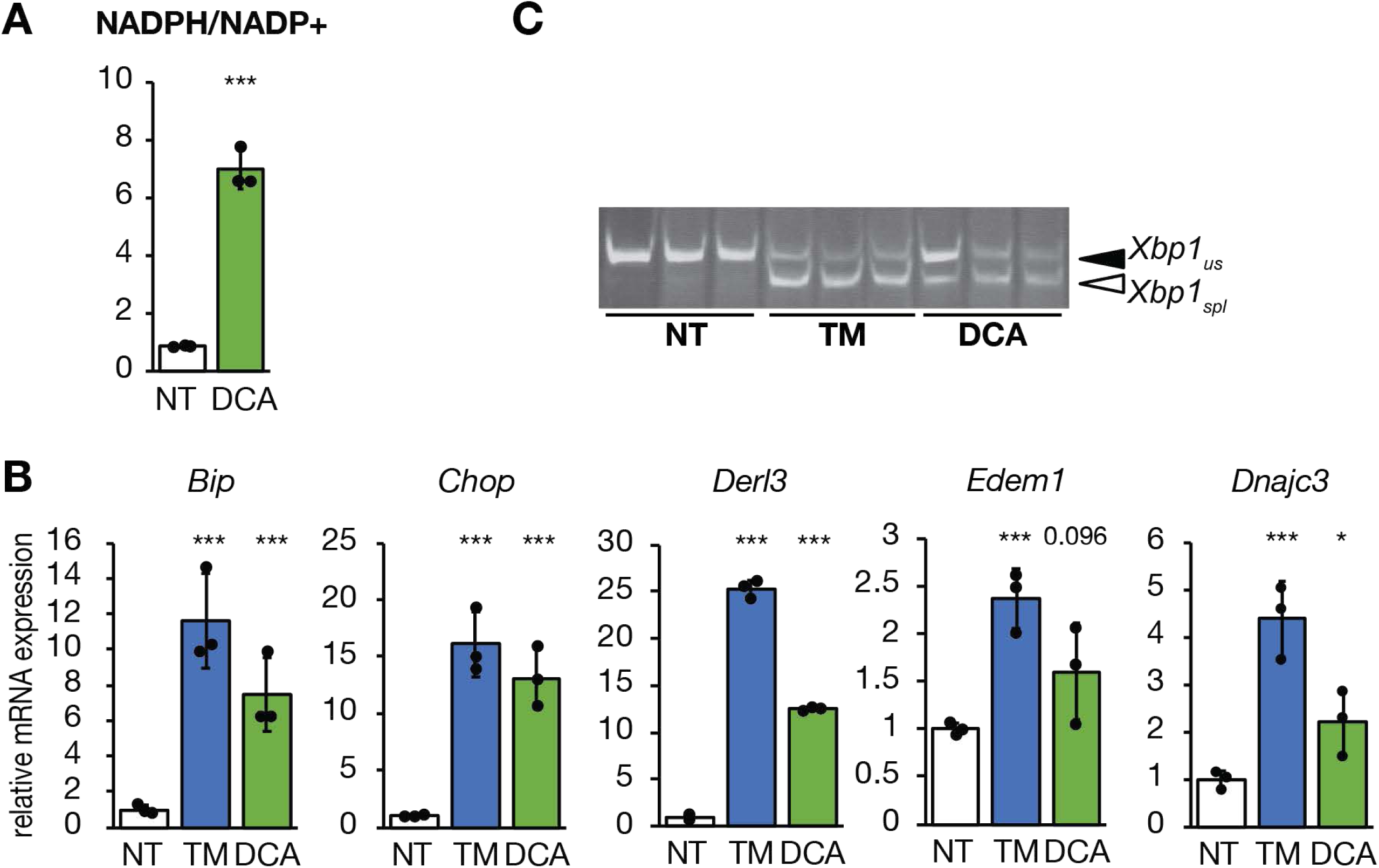
Stimulating TCA substrate entry causes ER stress. **(A-C)** Primary hepatocytes were treated with 4 µM dichloroacetate (DCA) or TM for 8 h. The NADPH/NADP+ ratio (A), UPR target gene expression (C), or *Xbp1* mRNA splicing (B) were analyzed as in previous figures.

### Ablation of the Mitochondrial Pyruvate Carrier alleviates ER stress

Finally, we wished to test whether ER stress could be alleviated by non-pharmacological manipulation of TCA cycle activity. The Mitochondrial Pyruvate Carrier (MPC) is composed of two subunits, MPC1 and MPC2, and mediates mitochondrial import of pyruvate (Bricker et al., 2012; Herzig et al., 2012). Loss of either subunit eliminates MPC activity, and liver-specific ablation of the MPC diminishes liver damage and inflammation in mice on obesogenic diets (Gray et al., 2015; McCommis et al., 2015; McCommis et al., 2017). In the absence of MPC activity, import of pyruvate into the mitochondria is greatly diminished, meaning that less acetyl-CoA can be produced by pyruvate dehydrogenase and therefore that TCA cycle flux is diminished (Gray et al., 2015; Rauckhorst et al., 2017). Therefore, ablating the MPC should mimic the effects of etomoxir treatment since both ultimately result in diminished acetyl-CoA availability for oxidation in the TCA cycle. To test this prediction, we first examined the effects of deleting MPC1 by CRISPR in C2C12 myocytes. We found that loss of MPC1 diminished the NADPH/NADP+ ratio and elevated the GSSG/GSH ratio in these cells (Figure 8A, B). Diminished UPR activation at both the protein (Figure 8C) and mRNA (Figure 8D) levels indicated that MPC1-deficient cells were also remarkably resistant to ER stress induced by TM or by the ER calcium-disrupting agent thapsigargin (TG). These findings were confirmed in primary hepatocytes taken from mice lacking MPC2 in the liver (*Mpc2^LKO^*), when compared to cells from mice with an intact allele (*Mpc2^fl/fl^*). In cells lacking MPC2, both the upregulation of UPR target genes (Figure 8E) and *Xbp1* mRNA splicing (Figure 8F, G) were attenuated. These findings provide direct genetic evidence that ER homeostasis is responsive to the availability of TCA cycle substrates.

**Figure 8:**
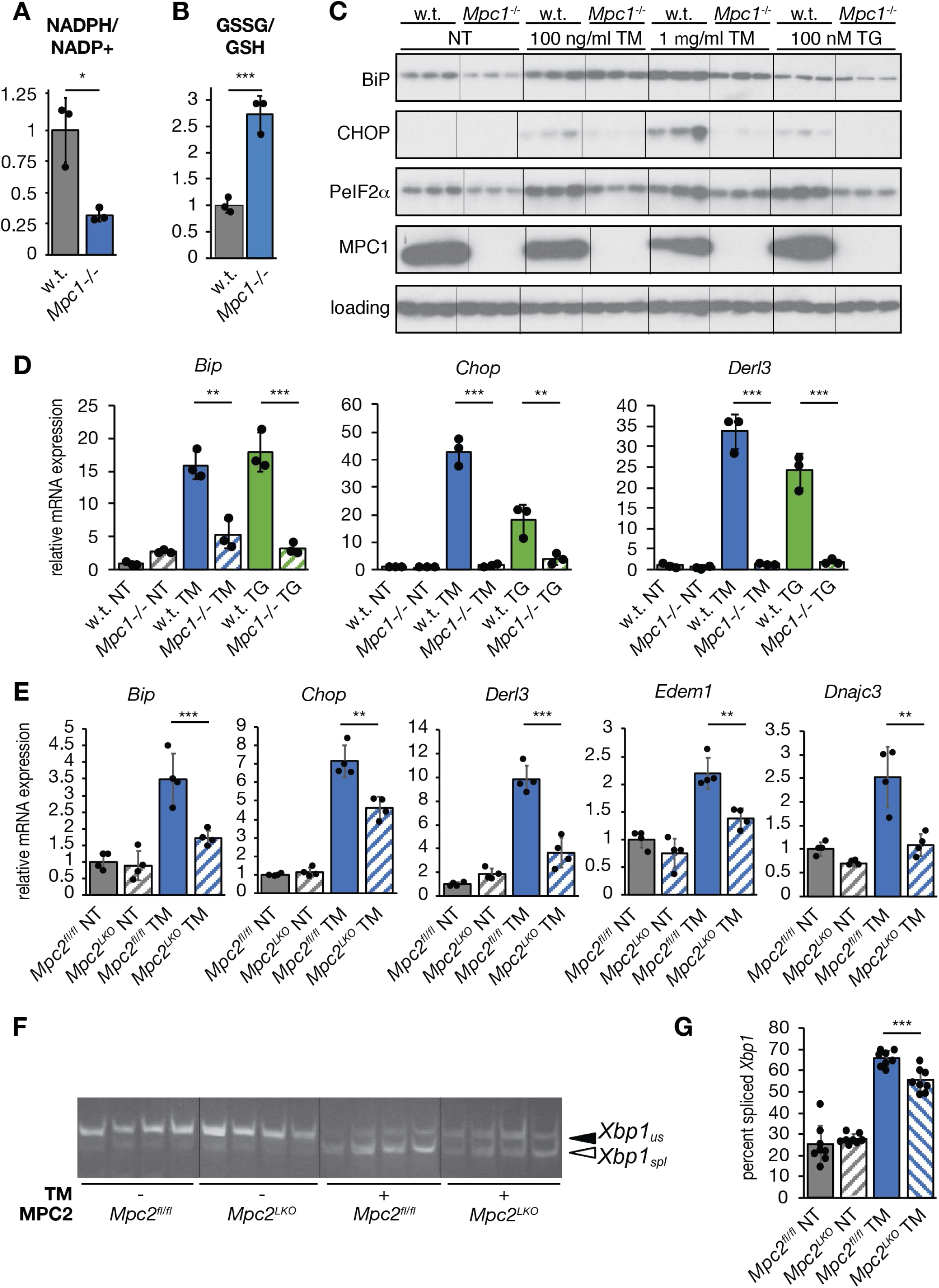
Attenuation of TCA flux by deletion of the Mitochondrial Pyruvate Carrier alleviates ER stress. **(A, B)** The NADPH/NADP+ ratio (A), and the GSSG/GSH ratio (B) measured as above in parental wild-type and *Mpc1*-/- C2C12 myoblasts. **(C)** Expression of BiP, CHOP, phosphorylated eIF2α, and MPC1 was analyzed by immunoblot in wild-type and *Mpc1*-/- C2C12 myocytes treated with 100 ng/mL TM, 1 μg/mL TM, or 100 nM thapsigargin (TG). Loading control was actin. **(D)** Relative mRNA expression of UPR markers from cells treated similarly to (C) was quantified by qRT-PCR. **(E)** Primary hepatocytes from *Mpc2^fl/fl^* and *Mpc2^LKO^* mice were treated with 250 ng/mL TM, and gene expression was measured by qRT-PCR. The *Mpc2*-/- untreated and w.t. TM-treated were reversed by cut/paste to keep ordering consistent with the rest of the data in the paper but the image was otherwise unmodified. **(F)** Splicing of *Xbp1* in TM-treated hepatocytes from *Mpc2^fl/fl^* or *Mpc2^LKO^*mice was assessed by conventional RT-PCR. (G) *Xbp1* splicing was quantified from 8 independent samples (4 from cells taken from a male mouse, 4 from cells taken from a female mouse).

## Discussion

Our results elucidate a pathway by which the ER senses metabolic activity (Figure 9). We propose that NADPH and GSSG convey TCA cycle status: decreasing the availability of acetyl-CoA—either from lipid or carbohydrate catabolism—dampens NADPH production and disfavors glutathione reduction, with the increased GSSG ultimately rendering the ER lumen more oxidizing and protecting ER homeostasis. Conversely, enhancing acetyl-CoA availability ultimately favors GSH over GSSG and causes ER stress. That these relationships were seen in hepatocytes, myocytes, and brown adipocytes, but were conspicuously absent in fibroblasts, hints that oxidative metabolism might represent an intrinsic challenge to ER functionality and might explain why feeding itself appears to be an ER stressor. Although there are many potential reasons that it might benefit the cell to tie ER protein processing to TCA cycle status— for example amino acid availability, ATP supply, lipid availability, etc.—the one we currently favor is the balancing of the cellular redox budget. Both the oxidation of nutrients in the mitochondria and the oxidation of nascent proteins in the ER generate toxic reactive oxygen species (ROS): in the mitochondria by electron leakage during oxidative phosphorylation (Murphy, 2009), and in the ER by catalysis of disulfide bond formation by ER oxidoreductase (ERO1) (Higa & Chevet, 2012). That inhibiting β-oxidation renders the ER more oxidizing suggests that the cell might be able to safely oxidize nutrients or nascent proteins, but not both simultaneously. As with the elucidation of any new pathway, our findings raise many further questions about how the pathway is initiated and regulated, and how it ultimately impacts cellular and organismal physiology.

**Figure 9:**
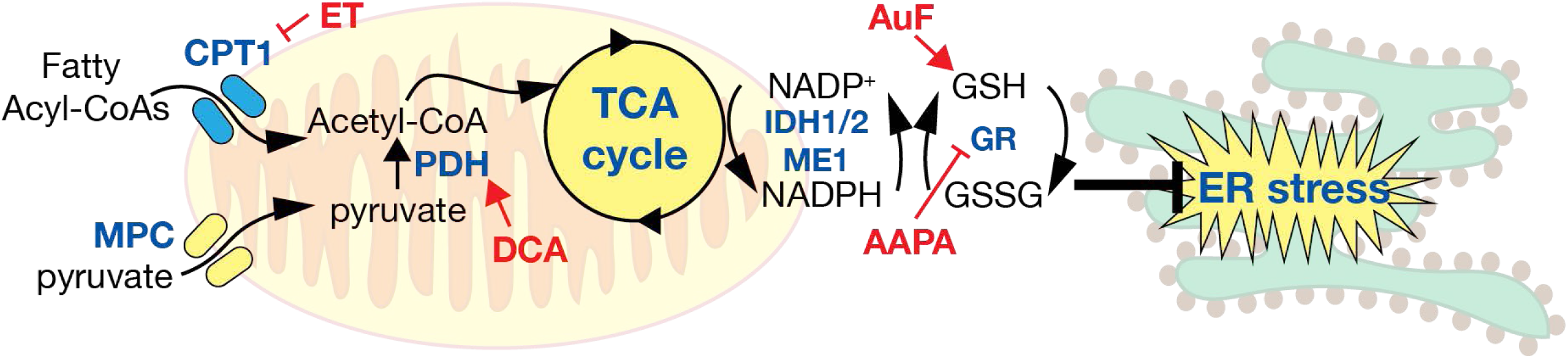
Proposed model describing the relationship between TCA cycle activity and ER homeostasis. See Discussion for details.

That increased TCA activity elicits ER stress does not necessarily portend cellular dysfunction. It is possible that ER stress induced by enhanced metabolic activity might “prime” the ER by eliciting prophylactic ER stress. Brief exposures to stress are known to precondition cells against subsequent stresses (Rutkowski & Kaufman, 2007). Metabolic fluxes are, at least outside the context of overnutrition and obesity, transient. Insulin action on the liver during feeding is known to promote both oxidative catabolism through PDH dephosphorylation (Moule & Denton, 1997; Wieland, Patzelt, & Loffler, 1972) and protein biogenesis through mTOR activation (Howell & Manning, 2011). UPR activation induced by enhanced TCA flux could enhance the functionality of the organelle in anticipation of the increase in protein synthesis— including synthesis of ER client proteins—that follows in the post-absorptive state.

While our data suggest that production of NADPH by the TCA cycle is a key event in our proposed pathway, it is not yet clear how NADPH production corresponds to actual activity of the cycle—given that metabolites can enter and exit the cycle at multiple points depending on nutritional conditions—nor how the cellular compartmentalization of NADPH production contributes to the transmission of the NADPH status to the ER. The NADPH/NADP+ ratio is regulated by nutritional state, exercise, diet, and circadian rhythms (Bradshaw, 2019; Ying, 2008). Only a portion of this NADPH is generated by the TCA cycle; other contributors include the pentose phosphate pathway of glucose metabolism and one-carbon folate metabolism, both of which are cytosolic. The fractional contribution of these pathways to total NADPH pools varies by cell type (J. Fan et al., 2014). In addition, we found here that inhibition of either the mitochondrial (IDH2) or cytosolic (IDH1 and ME1) NADPH-producing enzymes diminishes ER stress, and we have found previously that inhibition of the pentose phosphate pathway does likewise (Tyra et al., 2012). These findings imply that inhibiting NADPH production from any source protects ER function. How, then, are the cytosolic sources of NADPH linked to glutathione redox, since glutathione reductase contains a mitochondrial targeting signal? (Cytosolic glutathione reductase activity has been reported, possibly arising from translation initiation at a downstream start codon or inefficient mitochondrial targeting; (Kelner & Montoya, 2000)) Nicotinamide nucleotides are not thought to be competent for *trans*-mitochondrial transport (Lewis et al., 2014). One possibility is that diminishment of NADPH also inhibits thioredoxin reductase in the cytosol, and that cytosolic glutathione becomes more oxidized because it must compensate in the reduction of reactive oxygen species for the shortage of reduced thioredoxin. An alternate possibility is that cytosolic NADPH manipulation ultimately affects matrix NADPH status through metabolite shuttles, which are used to convey reducing equivalents across the mitochondrial membranes by coupling mitochondrial redox reactions with their cytosolic counterparts (for example, using the transport of citrate to couple IDH2 and IDH1) (Taylor, 2017). A next step will be to determine how TCA activity affects compartmentalization of NADPH arising from various sources.

Our results are surprising also in their implication that oxidized glutathione has a beneficial role in ER function. It is well-established that reducing agents elicit ER stress and activate the UPR, which speaks to the importance of the ER oxidative environment in the protein folding process. However, reducing equivalents are needed as well in order to activate ERO1 (Kim, Sideris, Sevier, & Kaiser, 2012) and to reduce disulfide isomerases so that they can catalyze reduction of improper disulfide bonds (Schwaller, Wilkinson, & Gilbert, 2003). Disruption of this capacity impairs the secretory pathway transport of model proteins with non-sequential disulfide bonds that must undergo such isomerization to avoid being trapped in non-native conformations (Poet et al., 2017). How oxidation and reduction are balanced within the ER lumen is not well-understood, and that understanding is confounded by the observations that neither ablation of both ERO1 isoforms (Zito, Chin, Blais, Harding, & Ron, 2010) nor apparent depletion of total ER glutathione (Tsunoda et al., 2014) appreciably impacts ER protein oxidation capacity or stress sensitivity except for specialized substrates. It could be that ER oxidative capacity and the relative need for oxidized glutathione varies by cell type.

An important question for future work is how elevated GSSG in the mitochondria and/or cytosol is transmitted to the ER. Does a change in the total cellular GSSG/GSH ratio ultimately change that ratio in the ER, or does GSSG exert its effects on the ER environment indirectly? To date, no mechanism for import of GSSG into the ER has been described. One possibility is that diminishing the cytosolic level of GSH suppresses its import into the ER, thereby elevating the ER GSSG/GSH ratio as well. However, given that the ratio of GSH to GSSG in the cytosol is approximately 100:1, it seems unlikely that cytosolic GSH is limiting in this way. Alternatively, perhaps sites of ER-mitochondrial contact facilitate spatially restricted exchange of glutathione as they do for other metabolites such as calcium and reactive oxygen species (Joseph, Booth, Young, & Hajnoczky, 2019). Ultimately, determining how elevated GSSG outside the ER leads to a more oxidizing ER lumen will require compartment-specific monitoring and manipulation of the glutathione redox state.

Our results also raise the question of what aspect(s) of ER functional capacity are altered by TCA cycle activity and glutathione redox. Activation of the UPR generally serves as a readout for ER stress, but it is not clear what diminished ER stress implies in terms of an actual accumulation of unfolded proteins, particularly because some stressors, such as palmitate loading, activate the UPR apparently independent of the protein folding process (Volmer et al., 2013). There are relatively few approaches for disentangling truly dysfunctional protein processing from the criteria by which the UPR perceives ER stress, which are still not well understood. For example, one might expect a dysfunctional ER to secrete proteins more slowly; on the other hand, perhaps enhanced ER retention time and chaperone association of client proteins facilitates proper protein folding under stressful conditions. In any case, we have yet to find any robust differences in nascent ER protein trafficking, processing, or aggregation in cells in which TCA activity is manipulated other than apparently enhanced clearance of NHK (Figure 4D). It remains to be determined whether this effect extends to other substrates and cell types. One intriguing—but speculative—possibility is that GSSG might paradoxically protect the ER by exacerbating protein misfolding, if doing so renders misfolded client proteins more readily recognized and either refolded or cleared rather than futilely engaged with the ER quality control machinery. Whatever the case, we infer that inhibiting TCA activity actually protects ER homeostasis in part because of its effects on ER ultrastructure. In principle, a hyperoxidizing ER might simply blunt the UPR rather than improve ER function, since activation of at least the UPR sensors ATF6 and IRE1 can be inhibited by oxidation (Nadanaka, Okada, Yoshida, & Mori, 2007; J. M. Wang et al., 2018). However, there is no evidence that enforcing reduction of these sensors is sufficient for their activation, meaning there would then be no reason to expect DCA to cause ER stress on its own. Moving forward, the redox status and activity of BiP will be of particular interest, given its roles in protein folding, ERAD, protein translocation, and UPR signaling (Pobre, Poet, & Hendershot, 2019), and the fact that oxidative modifications can alter the nature of its associations with nascent proteins (J. Wang, Pareja, Kaiser, & Sevier, 2014) and with the translocation channel (Ponsero et al., 2017). Perhaps manipulations of TCA activity will provide a new tool for understanding the relationships between ER oxidation, ER stress, and UPR activation.

Although our findings extend to other cell types beyond hepatocytes, they seem particularly relevant to liver disease. Obesity is the leading cause of non-alcoholic fatty liver disease (NAFLD) which, along with its downstream consequences—steatohepatitis, cirrhosis, and liver cancer—is the most common liver disease in the world (Araujo, Rosso, Bedogni, Tiribelli, & Bellentani, 2018). ER stress and dysregulation of the UPR are associated with NAFLD/NASH in humans (Gonzalez-Rodriguez et al., 2014; Lake et al., 2014; Lebeaupin et al., 2015; Lebeaupin et al., 2018), and in mice on NASH-promoting diets(Charlton et al., 2011; Rahman et al., 2007). ER stress might promote NASH by aggravating diet-induced steatosis(Lee, Scapa, Cohen, & Glimcher, 2008; Oyadomari et al., 2008; Rutkowski et al., 2008), and ER stress has also been shown to directly activate inflammatory signaling cascades(Lebeaupin et al., 2018; Özcan et al., 2004; Willy, Young, Stevens, Masuoka, & Wek, 2015) and to promote hepatocyte cell death(Iracheta-Vellve et al., 2016; Olivares & Henkel, 2015). NAFLD is associated with increased TCA flux in the liver (Rauckhorst et al., 2017; Satapati et al., 2012; Sunny, Parks, Browning, & Burgess, 2011), raising the question of whether NAFLD progression is accelerated by ER stress arising from this increased flux, and whether diminished hepatic TCA flux in mice lacking MPC activity protects the liver at least in part by alleviating or preventing ER stress. On one hand, oxidative stress is recognized to contribute to NAFLD progression, and glutathione is known to protect against NAFLD progression (Liu, Baker, Baker, & Zhu, 2015), although most of those studies have examined glutathione synthesis rather than glutathione redox. Conversely, our observation of elevated GSSG in cells lacking MPC1 (Figure 8B) suggests that, at a minimum, GSSG is not incompatible with diminished sensitivity to NAFLD. Whether TCA-dependent NADPH production contributes to NAFLD is not known. IDH1 and IDH2 function *in vivo* has mostly been examined in the context of transforming mutations in gliomas that change IDH activity. Both IDH1 and IDH2 are widely expressed, and to our knowledge no liver-specific deletion of either has been created. Thus, the role of the axis identified here in NAFLD has not been directly tested.

In conclusion, we have identified a novel NADPH- and glutathione-dependent pathway through which TCA cycle activity impacts ER homeostasis. We expect that this pathway will be relevant to the physiology and pathophysiology of liver, muscle, adipose, and other highly metabolically active cell types.

## Materials and Methods

### Cell culture and drug treatments

Primary hepatocytes were isolated from mice of both sexes between 6-12 weeks of age. Mice were anesthetized with isoflurane for the duration of the isolation. The liver was perfused through the portal vein with freshly prepared Perfusion Medium followed by digestion with Liver Digest Medium. Media formulas were as follows: Liver Perfusion Medium: HBSS, no calcium, no magnesium, no phenol red (Life Technologies, Carlsbad, CA), 0.5 mM EDTA, 0.5 mM EGTA, 25 mM HEPES, and penicillin-streptomycin (10,000 U/mL); Liver Digest Medium: HBSS, calcium, magnesium, no phenol red, 25 mM HEPES, penicillin-streptomycin (10,000 U/mL), 3.6 mg Trypsin Inhibitor (Sigma, St. Louis, MO), 25 mg Collagenase Type IV (210 U/mg) (Worthington Biochemical Corp., Lakewood, NJ). Flow rates were 4 ml/min for 5 min for perfusion, and 8 min for digestion. The liver was quickly excised, placed in cold Wash Medium (DMEM, 10 mM HEPES, 5% FBS, 100 μg/mL penicillin-streptomycin), dispersed by tearing Glisson’s capsule, and filtered through a sterile 70 μm cell strainer. Hepatocyte suspensions were centrifuged at 500 rpm for 3 min and resuspended in 30 mL of Wash Medium with 35% Percoll. Cells were centrifuged for 5 min at 1000 rpm, followed by resuspension in Wash Medium for a final wash with centrifugation for 3 min at 500 rpm. Viable hepatocytes were resuspended in Hepatocyte Medium (William’s E, 5% FBS, 10 nM insulin, 100 nM dexamethasone, 100 nM triiodothyronine, and 100 μg/mL penicillin-streptomycin), or, for *Mpc^fl/fl^*and liver-specific knockout (*Mpc2^LKO^*) hepatocytes, in high glucose DMEM, 10% FBS, penicillin-streptomycin, and 0.5 μg/ml amphotericin B, and plated on collagen-coated tissue culture plates. Media was changed 4 h after plating to remove any non-adherent cells. MPC2-deficient hepatocytes were isolated from *Mpc2^fl/fl^* animals bred into the Albumin-CRE line. MPC1-deficient C2C12 cells were generated by CRISPR and cultured as described (Oonthonpan, Rauckhorst, Gray, Boutron, & Taylor, 2019). gRNA sequences were 5′-GCGCTCCTACCGGTGCCCGA-3′ and 5′-GCCAACGGCACGGCCATGGC-3′.

Primary (pBAT) and immortalized (iBAT) brown adipocytes were isolated and cultured as described (Markan et al., 2014). Primary mouse embryonic fibroblasts were isolated and cultured as described (Scheuner et al., 2001). Drug treatments used the times and concentrations indicated in the figure legends. TM, auranofin, and DCA were from Millipore Sigma; etomoxir, 2-AAPA, and MG-132 from Cayman Chemical; stocks of each of these were stored at −20°C in DMSO. PA and OA (Millipore Sigma) were diluted stepwise in DMSO to 200 mM, and then to 100 mM in 10% fatty acid-free BSA (Millipore Sigma), followed by incubation at 40°C for 90 min with gentle agitation. Doxycycline (Millipore Sigma) stocks were stored at −20°C in water.

### Adenovirus experiments

A cDNA encoding the NHK allele of α1-antitrpysin was cloned into an adenoviral shuttle vector downstream of a TRE-Tight promoter, and the shuttle was recombined with a backbone expressing rtTA under the RSV promoter. Virus was purified by the University of Iowa Viral Vector Core. Primary hepatocytes were infected with Ad-TetOn-NHK or Ad-CMV-eGFP at a multiplicity of infection of 1:1. Infection began when media was changed 4 h after cells were plated. Cells were incubated with adenovirus for at least 12 h prior to addition of doxycycline or other treatments. NHK expression was induced with 500 ng/mL doxycycline in fresh media at the time of treatment.

### dsiRNA knockdown experiments

Primary hepatocytes were cultured overnight prior to transfection. Hepatocytes were transfected with 2.75 µM dsiRNA (Integrated DNA Technologies, Coralville, IA) in nuclease-free duplex buffer using the Viromer® Blue transfection kit (Origene) following the manufacturer’s protocol. A non-targeting dsiRNA was used as a control and two dsiRNAs for each gene of interest (*Idh1*, *Idh2*, and *Me1*) were used. Hepatocytes were transfected with dsiRNA for 24 h before experimental treatments. Targeting sequences were as follows: *Idh1*: GUACAACCAGGAUAAGUCAAUUGAA, GUUGAAGAAUUCAAGUUGAAACAAA; *Idh2*: AUUUAUAUUGCUCUGGAAUCACATG, AUCUUUGACAAGCACUAUAAGACTG; *Me1*: GCCAUUGUUCAAAAGAUAAAACCAA, ACCUUUCUAUCAGAUAUUAAAAUAT; non-targeting control: CGUUAAUCGCGUAUAAUACGCGUAT

### Biochemical Assays

Levels of total (GSx), oxidized (GSSG), and reduced (GSH) glutathione were measured using a Glutathione Fluorometric Assay Kit (Biovision, Milpitas, CA) following the manufacturer’s protocol. NADPH levels were measured using an NADP/NADPH Colorimetric Quantification Kit, and acetyl-CoA levels using the PicoProbe™ Acetyl-CoA Fluorometric Assay Kit (Biovision) following the manufacturer’s protocols, with the addition of 6N perchloric acid to precipitate proteins.

### RNA and Protein Analyses

Protein lysates were processed for immunoblot as described (Rutkowski et al. 2006). Primary antibodies were: CHOP (Santa Cruz sc-7351 or Proteintech 15204-1-AP), BiP (BD Biosciences 610978), PeIF2α (Invitrogen 44-728G), MPC1 (Cell Signaling Technology 14462), calnexin (loading control; Enzo ADI-SPA-865), actin (loading control; MP Biomedicals 691001), α1-antitrypsin (Dako A0012). The oxidative state of the ER was measured by incorporation of PEG-maleimide (mm(PEG)_24_) (ThermoFisher) as described (Tyra et al., 2012). Samples were run on Tris-tricine or Tris-HCl SDS-PAGE gels and transferred to 0.45 µm Immobilon-P Polyvinylidene Fluoride (PVDF) (Millipore) for Western blotting using ECL Prime substrate (GE Healthcare). qRT-PCR, including primer validation by standard curve and melt curve analysis, was as described (Rutkowski et al., 2006). Briefly, RNA was isolated following the standard Trizol protocol and RNA oncentrations were obtained using the Qubit RNA Broad Range kit. Concentrations were normalized, and cDNA was synthesized using 400 ng RNA with PrimeScript RT Master Mix (Takara). PCR reactions were performed using TB Green Premix Ex Taq (Takara) in a CFX96 cycler (Bio-Rad). Oligonucleotide sequences are listed in Table 1. Two loading controls were included in each experiment (*Btf3* and *Ppia*). Conventional RT-PCR was performed to assess splicing of *Xbp1* as described (Tyra et al., 2012).

**Table 1:**
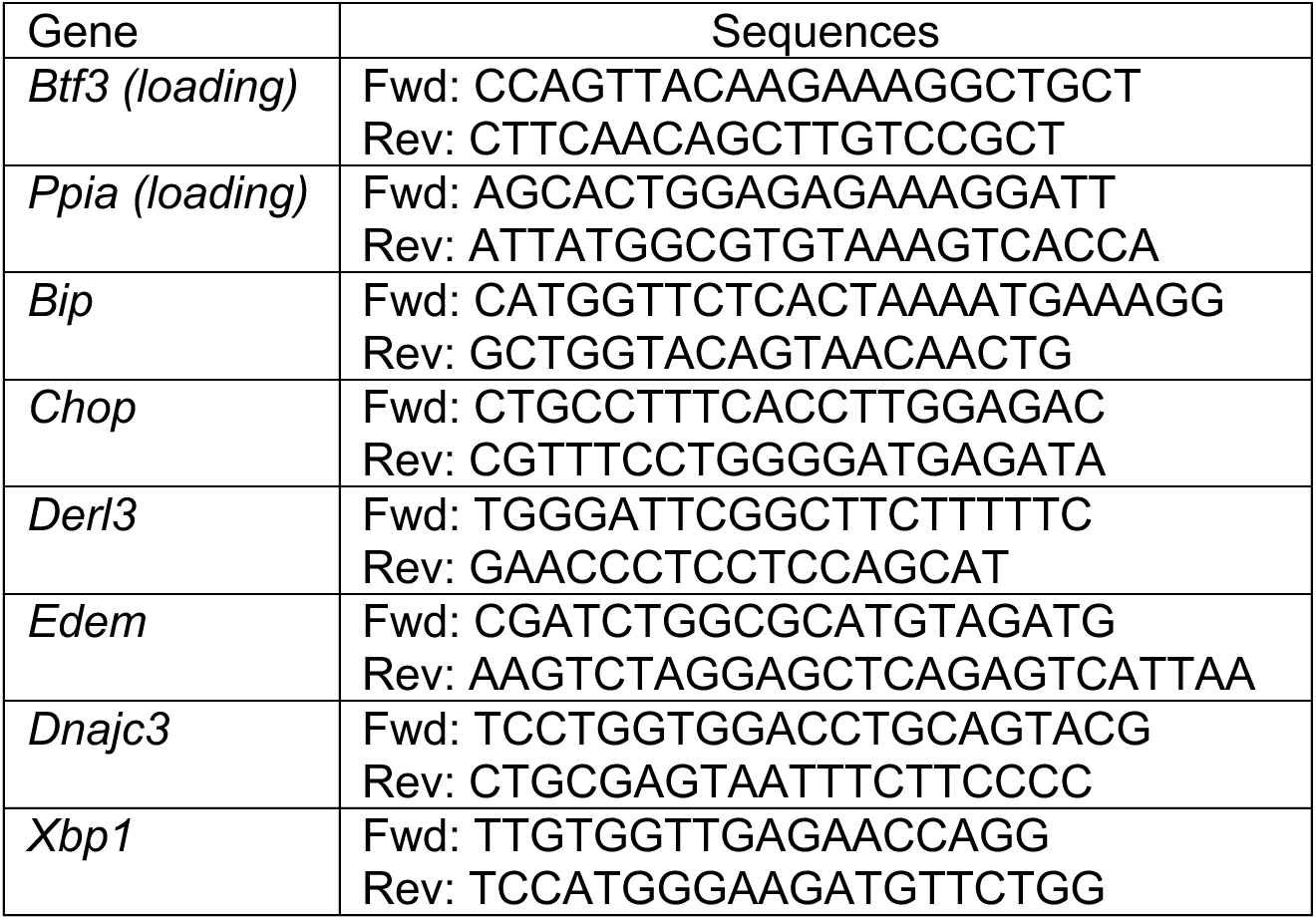
Oligonucleotide sequences.

### Transmission Electron Microscopy

Primary hepatocytes were cultured on collagen-coated glass coverslips and fixed in 2.5% glutaraldehyde prior to imaging by the University of Iowa Central Microscopy Research Facility. TEM images were scored blindly and binned into categories based on ER morphology: Normal (<25% dilated), Intermediate (25-75% dilated), or Severe (>75% dilated).

### Statistical Analyses

Continuous variables were reported as the mean ± standard deviation and were analyzed using the two-tailed Student’s t-test with Benjamini-Hochberg post-hoc correction for multiple comparisons. For qRT-PCR, significance was calculated prior to transformation of C_t_ values out of the log phase. A post-correction alpha of 0.05 was used to determine statistical significance.

## Acknowledgments and Funding

The authors would like to thank the University of Iowa Viral Vector Core and Central Microscopy Research Facility for technical assistance. Funding sources were as follows: DTR: R01GM115424 (NIH); ERG: T32GM067795 (NIH); MJP: R01DK106104 (NIH); BNF: DK104735 (NIH); EBT: DK104998 (NIH); and Central Microscopy Research Facility: 1 S10 RR018998 (NIH) (for JEOL JEM-1230 Transmission Electron Microscope)

## Author Contributions

ERG and DTR conceived and designed the experiments. ERG, KSM, MM, and AQK-M performed the experiments. ERG, KSM, MJP, BNF, EBT, and DTR interpreted the data. ERG and DTR wrote the manuscript. All authors edited and approved the manuscript.

---

**Figure S1.**
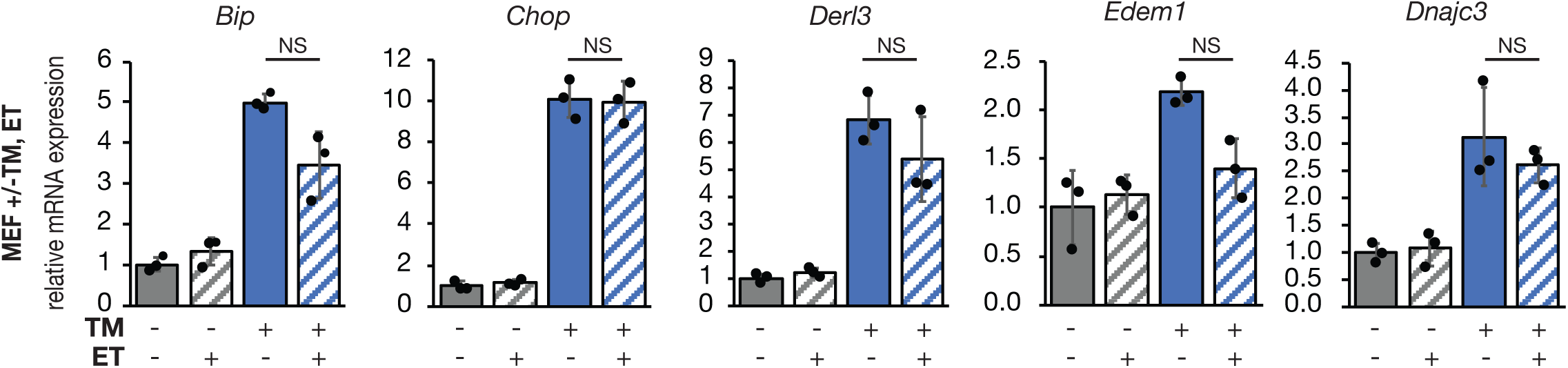
ET treatment does not diminish ER stress in primary mouse embryonic fibroblasts. Primary mouse embryonic fibroblasts treated were treated with 25 μg/ml ET and 250 ng/ml TM as indicated for 8h. mRNA expression was assessed by qRT-PCR as in Figure 1.

**Figure S2.**
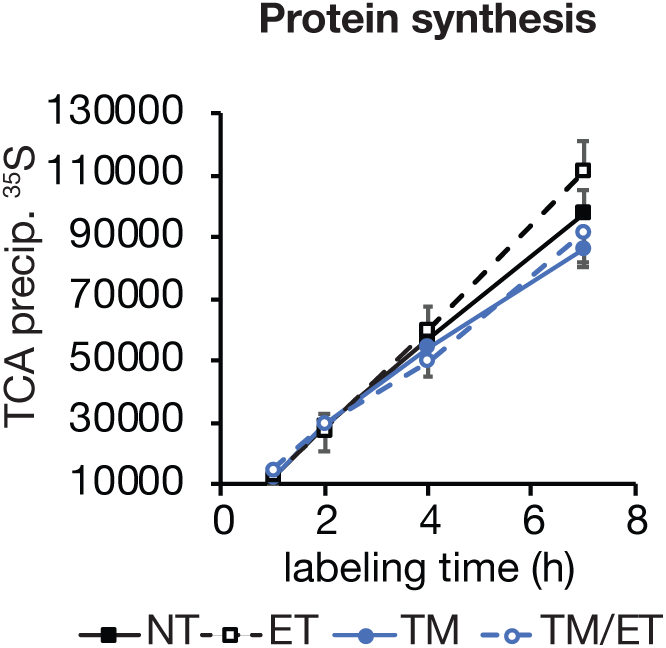
ET treatment does not affect protein synthesis. Primary hepatocytes treated were treated with 25 μg/ml ET and 250 ng/ml TM as indicated in media with 10 percent the normal amount of methionine and cysteine, and EasyTag ExpreSS^35^S (Perkin Elmer NEG772002MC) was added to a final concentration of 50 mCi/ml. After the indicated times, cell lysates were collected in 1% SDS 100 mM Tris, pH 8.8, denatured by heating, spotted onto gridded 3MM filter paper, and air dried. Filter was then precipitated by serial incubations in ice-cold 10% trichloroacetic acid (TCA) for 60 min., 5% TCA at room temperature for 5 min., 5% TCA preheated to 75° for 5 min., 5% TCA at room temperature for 5 min., and finally 2:1 ethanol:ether preheated to 37° for 5 min. Filter was then air dried, sliced into individual samples, and immersed in scintillation cocktail, and counted. n = 6 samples per condition. The modest effect of this dose of TM on protein synthesis is consistent with previous findings (Rutkowski et al., 2006)

## References

Araujo, A. R., Rosso, N., Bedogni, G., Tiribelli, C., & Bellentani, S. (2018). Global epidemiology of non-alcoholic fatty liver disease/non-alcoholic steatohepatitis: What we need in the future. Liver Int, 38 Suppl 1, 47–51. doi:10.1111/liv.13643

Bradshaw, P. C. (2019). Cytoplasmic and Mitochondrial NADPH-Coupled Redox Systems in the Regulation of Aging. Nutrients, 11(3). doi:10.3390/nu11030504

Bricker, D. K., Taylor, E. B., Schell, J. C., Orsak, T., Boutron, A., Chen, Y. C., & Rutter, J. (2012). A mitochondrial pyruvate carrier required for pyruvate uptake in yeast, Drosophila, and humans. Science, 337(6090), 96–100. doi:10.1126/science.1218099

Charlton, M., Krishnan, A., Viker, K., Sanderson, S., Cazanave, S., McConico, A., & Gores, G. (2011). Fast food diet mouse: novel small animal model of NASH with ballooning, progressive fibrosis, and high physiological fidelity to the human condition. Am J Physiol Gastrointest Liver Physiol, 301(5), G825–834. doi:10.1152/ajpgi.00145.2011

Chaudiere, J., & Tappel, A. L. (1984). Interaction of gold(I) with the active site of selenium-glutathione peroxidase. J Inorg Biochem, 20(4), 313–325. doi:10.1016/0162-0134(84)85030-8

Cnop, M., Foufelle, F., & Velloso, L. A. (2012). Endoplasmic reticulum stress, obesity and diabetes. Trends Mol Med, 18(1), 59–68. doi:10.1016/j.molmed.2011.07.010

Constantin-Teodosiu, D., Simpson, E. J., & Greenhaff, P. L. (1999). The importance of pyruvate availability to PDC activation and anaplerosis in human skeletal muscle. Am J Physiol, 276(3), E472–478. doi:10.1152/ajpendo.1999.276.3.E472

Delaunay-Moisan, A., Ponsero, A., & Toledano, M. B. (2017). Reexamining the Function of Glutathione in Oxidative Protein Folding and Secretion. Antioxid Redox Signal, 27(15), 1178–1199. doi:10.1089/ars.2017.7148

DeZwaan-McCabe, D., Sheldon, R. D., Gorecki, M. C., Guo, D. F., Gansemer, E. R., Kaufman, R. J., & Rutkowski, D. T. (2017). ER Stress Inhibits Liver Fatty Acid Oxidation while Unmitigated Stress Leads to Anorexia-Induced Lipolysis and Both Liver and Kidney Steatosis. Cell Rep, 19(9), 1794–1806. doi:10.1016/j.celrep.2017.05.020

Fan, J., Ye, J., Kamphorst, J. J., Shlomi, T., Thompson, C. B., & Rabinowitz, J. D. (2014). Quantitative flux analysis reveals folate-dependent NADPH production. Nature, 510(7504), 298–302. doi:10.1038/nature13236

Fan, Y., & Simmen, T. (2019). Mechanistic Connections between Endoplasmic Reticulum (ER) Redox Control and Mitochondrial Metabolism. Cells, 8(9). doi:10.3390/cells8091071

Finnie, J. W. (2001). Effect of tunicamycin on hepatocytes in vitro. J Comp Pathol, 125(4), 318–321. doi:10.1053/jcpa.2001.0510

Frand, A. R., & Kaiser, C. A. (1999). Ero1p oxidizes protein disulfide isomerase in a pathway for disulfide bond formation in the endoplasmic reticulum. Mol Cell, 4(4), 469–477. Retrieved from http://www.ncbi.nlm.nih.gov/pubmed/10549279

Frayn, K. N., Arner, P., & Yki-Jarvinen, H. (2006). Fatty acid metabolism in adipose tissue, muscle and liver in health and disease. Essays Biochem, 42, 89–103. doi:10.1042/bse0420089

Fu, S., Watkins, S. M., & Hotamisligil, G. S. (2012). The role of endoplasmic reticulum in hepatic lipid homeostasis and stress signaling. Cell Metab, 15(5), 623–634. doi:10.1016/j.cmet.2012.03.007

Gomez, J. A., & Rutkowski, D. T. (2016). Experimental reconstitution of chronic ER stress in the liver reveals feedback suppression of BiP mRNA expression. Elife, 5. doi:10.7554/eLife.20390

Gonzalez-Rodriguez, A., Mayoral, R., Agra, N., Valdecantos, M. P., Pardo, V., Miquilena-Colina, M. E., & Valverde, A. M. (2014). Impaired autophagic flux is associated with increased endoplasmic reticulum stress during the development of NAFLD. Cell Death Dis, 5, e1179. doi:10.1038/cddis.2014.162

Gray, L. R., Sultana, M. R., Rauckhorst, A. J., Oonthonpan, L., Tompkins, S. C., Sharma, A., & Taylor, E. B. (2015). Hepatic Mitochondrial Pyruvate Carrier 1 Is Required for Efficient Regulation of Gluconeogenesis and Whole-Body Glucose Homeostasis. Cell Metab, 22(4), 669–681. doi:10.1016/j.cmet.2015.07.027

Gray, L. R., Tompkins, S. C., & Taylor, E. B. (2014). Regulation of pyruvate metabolism and human disease. Cell Mol Life Sci, 71(14), 2577–2604. doi:10.1007/s00018-013-1539-2

Gutierrez, T., & Simmen, T. (2018). Endoplasmic reticulum chaperones tweak the mitochondrial calcium rheostat to control metabolism and cell death. Cell Calcium, 70, 64–75. doi:10.1016/j.ceca.2017.05.015

Herzig, S., Raemy, E., Montessuit, S., Veuthey, J. L., Zamboni, N., Westermann, B., & Martinou, J. C. (2012). Identification and functional expression of the mitochondrial pyruvate carrier. Science, 337(6090), 93–96. doi:10.1126/science.1218530

Higa, A., & Chevet, E. (2012). Redox signaling loops in the unfolded protein response. Cell Signal, 24(8), 1548–1555. doi:10.1016/j.cellsig.2012.03.011

Howell, J. J., & Manning, B. D. (2011). mTOR couples cellular nutrient sensing to organismal metabolic homeostasis. Trends Endocrinol Metab, 22(3), 94–102. doi:10.1016/j.tem.2010.12.003

Iracheta-Vellve, A., Petrasek, J., Gyongyosi, B., Satishchandran, A., Lowe, P., Kodys, K., & Szabo, G. (2016). Endoplasmic Reticulum Stress-induced Hepatocellular Death Pathways Mediate Liver Injury and Fibrosis via Stimulator of Interferon Genes. J Biol Chem, 291(52), 26794–26805. doi:10.1074/jbc.M116.736991

Joseph, S. K., Booth, D. M., Young, M. P., & Hajnoczky, G. (2019). Redox regulation of ER and mitochondrial Ca(2+) signaling in cell survival and death. Cell Calcium, 79, 89–97. doi:10.1016/j.ceca.2019.02.006

Kelner, M. J., & Montoya, M. A. (2000). Structural organization of the human glutathione reductase gene: determination of correct cDNA sequence and identification of a mitochondrial leader sequence. Biochem Biophys Res Commun, 269(2), 366–368. doi:10.1006/bbrc.2000.2267

Kim, S., Sideris, D. P., Sevier, C. S., & Kaiser, C. A. (2012). Balanced Ero1 activation and inactivation establishes ER redox homeostasis. J Cell Biol, 196(6), 713–725. doi:10.1083/jcb.201110090

Lai, K. K., Kolippakkam, D., & Beretta, L. (2008). Comprehensive and quantitative proteome profiling of the mouse liver and plasma. Hepatology, 47(3), 1043–1051. doi:10.1002/hep.22123

Lake, A. D., Novak, P., Hardwick, R. N., Flores-Keown, B., Zhao, F., Klimecki, W. T., & Cherrington, N. J. (2014). The adaptive endoplasmic reticulum stress response to lipotoxicity in progressive human nonalcoholic fatty liver disease. Toxicol Sci, 137(1), 26–35. doi:10.1093/toxsci/kft230

Lebeaupin, C., Proics, E., de Bieville, C. H., Rousseau, D., Bonnafous, S., Patouraux, S., & Bailly-Maitre, B. (2015). ER stress induces NLRP3 inflammasome activation and hepatocyte death. Cell Death Dis, 6, e1879. doi:10.1038/cddis.2015.248

Lebeaupin, C., Vallee, D., Rousseau, D., Patouraux, S., Bonnafous, S., Adam, G., & Bailly-Maitre, B. (2018). Bax inhibitor-1 protects from nonalcoholic steatohepatitis by limiting inositol-requiring enzyme 1 alpha signaling in mice. Hepatology, 68(2), 515–532. doi:10.1002/hep.29847

Lee, A. H., Scapa, E. F., Cohen, D. E., & Glimcher, L. H. (2008). Regulation of hepatic lipogenesis by the transcription factor XBP1. Science, 320(5882), 1492–1496. doi:10.1126/science.1158042

Lewis, C. A., Parker, S. J., Fiske, B. P., McCloskey, D., Gui, D. Y., Green, C. R., & Metallo, C. (2014). Tracing compartmentalized NADPH metabolism in the cytosol and mitochondria of mammalian cells. Mol Cell, 55(2), 253–263. doi:10.1016/j.molcel.2014.05.008

Liu, W., Baker, S. S., Baker, R. D., & Zhu, L. (2015). Antioxidant Mechanisms in Nonalcoholic Fatty Liver Disease. Curr Drug Targets, 16(12), 1301–1314. doi:10.2174/1389450116666150427155342

Markan, K. R., Naber, M. C., Ameka, M. K., Anderegg, M. D., Mangelsdorf, D. J., Kliewer, S. A., & Potthoff, M. J. (2014). Circulating FGF21 is liver derived and enhances glucose uptake during refeeding and overfeeding. Diabetes, 63(12), 4057–4063. doi:10.2337/db14-0595

McCommis, K. S., Chen, Z., Fu, X., McDonald, W. G., Colca, J. R., Kletzien, R. F., & Finck, B. (2015). Loss of Mitochondrial Pyruvate Carrier 2 in the Liver Leads to Defects in Gluconeogenesis and Compensation via Pyruvate-Alanine Cycling. Cell Metab, 22(4), 682–694. doi:10.1016/j.cmet.2015.07.028

McCommis, K. S., Hodges, W. T., Brunt, E. M., Nalbantoglu, I., McDonald, W. G., Holley, C., & Finck, B. N. (2017). Targeting the mitochondrial pyruvate carrier attenuates fibrosis in a mouse model of nonalcoholic steatohepatitis. Hepatology, 65(5), 1543–1556. doi:10.1002/hep.29025

Merrill, C. L., Ni, H., Yoon, L. W., Tirmenstein, M. A., Narayanan, P., Benavides, G. R., & Morgan, K. T. (2002). Etomoxir-induced oxidative stress in HepG2 cells detected by differential gene expression is confirmed biochemically. Toxicol Sci, 68(1), 93–101. Retrieved from http://www.ncbi.nlm.nih.gov/entrez/query.fcgi?cmd=Retrieve&db=PubMed&dopt=Citation&list_uids=12075114

Mogilenko, D. A., Haas, J. T., L’Homme, L., Fleury, S., Quemener, S., Levavasseur, M., & Dombrowicz, D. (2019). Metabolic and Innate Immune Cues Merge into a Specific Inflammatory Response via the UPR. Cell, 178(1), 263. doi:10.1016/j.cell.2019.06.017

Mohan, S., R, P. R. M., Brown, L., Ayyappan, P., & G, R. K. (2019). Endoplasmic reticulum stress: A master regulator of metabolic syndrome. Eur J Pharmacol, 860, 172553. doi:10.1016/j.ejphar.2019.172553

Moule, S. K., & Denton, R. M. (1997). Multiple signaling pathways involved in the metabolic effects of insulin. Am J Cardiol, 80(3A), 41A–49A. doi:10.1016/s0002-9149(97)00457-8

Murphy, M. P. (2009). How mitochondria produce reactive oxygen species. Biochem J, 417(1), 1–13. doi:10.1042/BJ20081386

Nadanaka, S., Okada, T., Yoshida, H., & Mori, K. (2007). Role of disulfide bridges formed in the luminal domain of ATF6 in sensing endoplasmic reticulum stress. Mol Cell Biol, 27(3), 1027–1043. doi:MCB.00408-06 [pii]10.1128/MCB.00408-06

Nagasawa, K., Higashi, T., Hosokawa, N., Kaufman, R. J., & Nagata, K. (2007). Simultaneous induction of the four subunits of the TRAP complex by ER stress accelerates ER degradation. EMBO Rep, 8(5), 483–489. Retrieved from http://www.ncbi.nlm.nih.gov/entrez/query.fcgi?cmd=Retrieve&db=PubMed&dopt=Citation&list_uids=17380188

Olivares, S., & Henkel, A. S. (2015). Hepatic Xbp1 Gene Deletion Promotes Endoplasmic Reticulum Stress-induced Liver Injury and Apoptosis. J Biol Chem, 290(50), 30142–30151. doi:10.1074/jbc.M115.676239

Oonthonpan, L., Rauckhorst, A. J., Gray, L. R., Boutron, A. C., & Taylor, E. B. (2019). Two human patient mitochondrial pyruvate carrier mutations reveal distinct molecular mechanisms of dysfunction. JCI Insight, 5. doi:10.1172/jci.insight.126132

Ordonez, A., Snapp, E. L., Tan, L., Miranda, E., Marciniak, S. J., & Lomas, D. A. (2013). Endoplasmic reticulum polymers impair luminal protein mobility and sensitize to cellular stress in alpha1-antitrypsin deficiency. Hepatology, 57(5), 2049–2060. doi:10.1002/hep.26173

Owen, O. E., Kalhan, S. C., & Hanson, R. W. (2002). The key role of anaplerosis and cataplerosis for citric acid cycle function. J Biol Chem, 277(34), 30409–30412. doi:10.1074/jbc.R200006200

Oyadomari, S., Harding, H. P., Zhang, Y., Oyadomari, M., & Ron, D. (2008). Dephosphorylation of translation initiation factor 2alpha enhances glucose tolerance and attenuates hepatosteatosis in mice. Cell Metab, 7(6), 520–532.

Özcan, U., Cao, Q., Yilmaz, E., Lee, A. H., Iwakoshi, N. N., Ozdelen, E., & Hotamisligil, G. S. (2004). Endoplasmic reticulum stress links obesity, insulin action, and type 2 diabetes. Science, 306(5695), 457–461. Retrieved from http://www.ncbi.nlm.nih.gov/entrez/query.fcgi?cmd=Retrieve&db=PubMed&dopt=Citation&list_uids=15486293

Peters, T., Jr., & Davidson, L. K. (1982). The biosynthesis of rat serum albumin. In vivo studies on the formation of the disulfide bonds. J Biol Chem, 257(15), 8847–8853. Retrieved from https://www.ncbi.nlm.nih.gov/pubmed/7096338

Pfaffenbach, K. T., Nivala, A. M., Reese, L., Ellis, F., Wang, D., Wei, Y., & Pagliassotti, M. J. (2010). Rapamycin Inhibits Postprandial-Mediated X-Box-Binding Protein-1 Splicing in Rat Liver. J Nutr, 140(5), 879–884. doi:jn.109.119883 [pii]10.3945/jn.109.119883

Pike, L. S., Smift, A. L., Croteau, N. J., Ferrick, D. A., & Wu, M. (2010). Inhibition of fatty acid oxidation by etomoxir impairs NADPH production and increases reactive oxygen species resulting in ATP depletion and cell death in human glioblastoma cells. Biochim Biophys Acta, 1807(6), 726–734. Retrieved from http://www.ncbi.nlm.nih.gov/entrez/query.fcgi?cmd=Retrieve&db=PubMed&dopt=Citation&list_uids=21692241

Pobre, K. F. R., Poet, G. J., & Hendershot, L. M. (2019). The endoplasmic reticulum (ER) chaperone BiP is a master regulator of ER functions: Getting by with a little help from ERdj friends. J Biol Chem, 294(6), 2098–2108. doi:10.1074/jbc.REV118.002804

Poet, G. J., Oka, O. B., van Lith, M., Cao, Z., Robinson, P. J., Pringle, M. A., & Bulleid, N. J. (2017). Cytosolic thioredoxin reductase 1 is required for correct disulfide formation in the ER. Embo J, 36(5), 693–702. doi:10.15252/embj.201695336

Ponsero, A. J., Igbaria, A., Darch, M. A., Miled, S., Outten, C. E., Winther, J. R., & Toledano, M. B. (2017). Endoplasmic Reticulum Transport of Glutathione by Sec61 Is Regulated by Ero1 and Bip. Mol Cell, 67(6), 962–973 e965. doi:10.1016/j.molcel.2017.08.012

Raffaello, A., Mammucari, C., Gherardi, G., & Rizzuto, R. (2016). Calcium at the Center of Cell Signaling: Interplay between Endoplasmic Reticulum, Mitochondria, and Lysosomes. Trends Biochem Sci, 41(12), 1035–1049. doi:10.1016/j.tibs.2016.09.001

Rahman, S. M., Schroeder-Gloeckler, J. M., Janssen, R. C., Jiang, H., Qadri, I., Maclean, K. N., & Friedman, J. E. (2007). CCAAT/enhancing binding protein beta deletion in mice attenuates inflammation, endoplasmic reticulum stress, and lipid accumulation in diet-induced nonalcoholic steatohepatitis. Hepatology, 45(5), 1108–1117. doi:10.1002/hep.21614

Rainbolt, T. K., Saunders, J. M., & Wiseman, R. L. (2014). Stress-responsive regulation of mitochondria through the ER unfolded protein response. Trends Endocrinol Metab, 25(10), 528–537. doi:10.1016/j.tem.2014.06.007

Rauckhorst, A. J., Gray, L. R., Sheldon, R. D., Fu, X., Pewa, A. D., Feddersen, C. R., & Taylor, E. B. (2017). The mitochondrial pyruvate carrier mediates high fat diet-induced increases in hepatic TCA cycle capacity. Mol Metab, 6(11), 1468–1479. doi:10.1016/j.molmet.2017.09.002

Roberts, J. R., & Shaw, C. F., 3rd. (1998). Inhibition of erythrocyte selenium-glutathione peroxidase by auranofin analogues and metabolites. Biochem Pharmacol, 55(8), 1291–1299. doi:10.1016/s0006-2952(97)00634-5

Rutkowski, D. T., Arnold, S. M., Miller, C. N., Wu, J., Li, J., Gunnison, K. M., & Kaufman, R. J. (2006). Adaptation to ER stress is mediated by differential stabilities of pro-survival and pro-apoptotic mRNAs and proteins. PLoS Biol, 4(11), e374. doi:06-PLBI-RA-0091R3 [pii] 10.1371/journal.pbio.0040374

Rutkowski, D. T., & Kaufman, R. J. (2007). That which does not kill me makes me stronger: adapting to chronic ER stress. Trends Biochem Sci, 32(10), 469–476. doi:S0968-0004(07)00213-7 [pii]10.1016/j.tibs.2007.09.003

Rutkowski, D. T., Wu, J., Back, S. H., Callaghan, M. U., Ferris, S. P., Iqbal, J., & Kaufman, R. J. (2008). UPR pathways combine to prevent hepatic steatosis caused by ER stress-mediated suppression of transcriptional master regulators. Dev Cell, 15(6), 829–840. doi:S1534-5807(08)00475-9 [pii]10.1016/j.devcel.2008.10.015

Rydstrom, J. (2006). Mitochondrial NADPH, transhydrogenase and disease. Biochim Biophys Acta, 1757(5-6), 721–726. doi:10.1016/j.bbabio.2006.03.010

Salvado, L., Palomer, X., Barroso, E., & Vazquez-Carrera, M. (2015). Targeting endoplasmic reticulum stress in insulin resistance. Trends Endocrinol Metab, 26(8), 438–448. doi:10.1016/j.tem.2015.05.007

Satapati, S., Sunny, N. E., Kucejova, B., Fu, X., He, T. T., Mendez-Lucas, A., & Burgess, S. C. (2012). Elevated TCA cycle function in the pathology of diet-induced hepatic insulin resistance and fatty liver. J Lipid Res, 53(6), 1080–1092. doi:10.1194/jlr.M023382

Scarbrough, P. M., Mapuskar, K. A., Mattson, D. M., Gius, D., Watson, W. H., & Spitz, D. R. (2012). Simultaneous inhibition of glutathione- and thioredoxin-dependent metabolism is necessary to potentiate 17AAG-induced cancer cell killing via oxidative stress. Free Radic Biol Med, 52(2), 436–443. doi:10.1016/j.freeradbiomed.2011.10.493

Scheuner, D., Song, B., McEwen, E., Liu, C., Laybutt, R., Gillespie, P., & Kaufman, R. J. (2001). Translational control is required for the unfolded protein response and in vivo glucose homeostasis. Mol Cell, 7(6), 1165–1176. Retrieved from http://www.ncbi.nlm.nih.gov/entrez/query.fcgi?cmd=Retrieve&db=PubMed&dopt=Citation&list_uids=11430820

Schwaller, M., Wilkinson, B., & Gilbert, H. F. (2003). Reduction-reoxidation cycles contribute to catalysis of disulfide isomerization by protein-disulfide isomerase. J Biol Chem, 278(9), 7154–7159. doi:10.1074/jbc.M211036200

Seefeldt, T., Zhao, Y., Chen, W., Raza, A. S., Carlson, L., Herman, J., & Guan, X. (2009). Characterization of a novel dithiocarbamate glutathione reductase inhibitor and its use as a tool to modulate intracellular glutathione. J Biol Chem, 284(5), 2729–2737. doi:10.1074/jbc.M802683200

Sies, H., Berndt, C., & Jones, D. P. (2017). Oxidative Stress. Annu Rev Biochem, 86, 715–748. doi:10.1146/annurev-biochem-061516-045037

Sifers, R. N., Brashears-Macatee, S., Kidd, V. J., Muensch, H., & Woo, S. L. (1988). A frameshift mutation results in a truncated alpha 1-antitrypsin that is retained within the rough endoplasmic reticulum. J Biol Chem, 263(15), 7330–7335.

Sunny, N. E., Parks, E. J., Browning, J. D., & Burgess, S. C. (2011). Excessive hepatic mitochondrial TCA cycle and gluconeogenesis in humans with nonalcoholic fatty liver disease. Cell Metab, 14(6), 804–810. doi:10.1016/j.cmet.2011.11.004

Taylor, E. B. (2017). Functional Properties of the Mitochondrial Carrier System. Trends Cell Biol, 27(9), 633–644. doi:10.1016/j.tcb.2017.04.004

Theurey, P., Tubbs, E., Vial, G., Jacquemetton, J., Bendridi, N., Chauvin, M. A., & Rieusset, J. (2016). Mitochondria-associated endoplasmic reticulum membranes allow adaptation of mitochondrial metabolism to glucose availability in the liver. J Mol Cell Biol, 8(2), 129–143. doi:10.1093/jmcb/mjw004

Tsunoda, S., Avezov, E., Zyryanova, A., Konno, T., Mendes-Silva, L., Pinho Melo, E., & Ron, D. (2014). Intact protein folding in the glutathione-depleted endoplasmic reticulum implicates alternative protein thiol reductants. Elife, 3, e03421. doi:10.7554/eLife.03421

Tu, B. P., Ho-Schleyer, S. C., Travers, K. J., & Weissman, J. S. (2000). Biochemical basis of oxidative protein folding in the endoplasmic reticulum. Science, 290(5496), 1571–1574. Retrieved from http://www.ncbi.nlm.nih.gov/pubmed/11090354

Tyra, H. M., Spitz, D. R., & Rutkowski, D. T. (2012). Inhibition of fatty acid oxidation enhances oxidative protein folding and protects hepatocytes from endoplasmic reticulum stress. Mol Biol Cell, 23(5), 811–819. doi:mbc.E11-12-1011 [pii]10.1091/mbc.E11-12-1011

Vance, J. E. (2014). MAM (mitochondria-associated membranes) in mammalian cells: lipids and beyond. Biochim Biophys Acta, 1841(4), 595–609. doi:10.1016/j.bbalip.2013.11.014

Volmer, R., van der Ploeg, K., & Ron, D. (2013). Membrane lipid saturation activates endoplasmic reticulum unfolded protein response transducers through their transmembrane domains. Proc Natl Acad Sci U S A, 110(12), 4628–4633. doi:10.1073/pnas.1217611110

Walter, P., & Ron, D. (2011). The unfolded protein response: from stress pathway to homeostatic regulation. Science, 334(6059), 1081–1086. doi:334/6059/1081 [pii]10.1126/science.1209038

Wang, J., Pareja, K. A., Kaiser, C. A., & Sevier, C. S. (2014). Redox signaling via the molecular chaperone BiP protects cells against endoplasmic reticulum-derived oxidative stress. Elife, 3, e03496. doi:10.7554/eLife.03496

Wang, J. M., Qiu, Y., Yang, Z., Kim, H., Qian, Q., Sun, Q., & Zhang, K. (2018). IRE1alpha prevents hepatic steatosis by processing and promoting the degradation of select microRNAs. Sci Signal, 11(530). doi:10.1126/scisignal.aao4617

Weis, B. C., Cowan, A. T., Brown, N., Foster, D. W., & McGarry, J. D. (1994). Use of a selective inhibitor of liver carnitine palmitoyltransferase I (CPT I) allows quantification of its contribution to total CPT I activity in rat heart. Evidence that the dominant cardiac CPT I isoform is identical to the skeletal muscle enzyme. J Biol Chem, 269(42), 26443–26448. Retrieved from http://www.ncbi.nlm.nih.gov/entrez/query.fcgi?cmd=Retrieve&db=PubMed&dopt=Citation&list_uids=7929365

Wieland, O. H., Patzelt, C., & Loffler, G. (1972). Active and inactive forms of pyruvate dehydrogenase in rat liver. Effect of starvation and refeeding and of insulin treatment on pyruvate-dehydrogenase interconversion. Eur J Biochem, 26(3), 426–433. doi:10.1111/j.1432-1033.1972.tb01783.x

Willy, J. A., Young, S. K., Stevens, J. L., Masuoka, H. C., & Wek, R. C. (2015). CHOP links endoplasmic reticulum stress to NF-kappaB activation in the pathogenesis of nonalcoholic steatohepatitis. Mol Biol Cell, 26(12), 2190–2204. doi:10.1091/mbc.E15-01-0036

Winther, J. R., & Thorpe, C. (2014). Quantification of thiols and disulfides. Biochim Biophys Acta, 1840(2), 838–846. doi:10.1016/j.bbagen.2013.03.031

Wu, C. Y., Satapati, S., Gui, W., Wynn, R. M., Sharma, G., Lou, M., & Merritt, M. E. (2018). A novel inhibitor of pyruvate dehydrogenase kinase stimulates myocardial carbohydrate oxidation in diet-induced obesity. J Biol Chem, 293(25), 9604–9613. doi:10.1074/jbc.RA118.002838

Wu, J., Rutkowski, D. T., Dubois, M., Swathirajan, J., Saunders, T., Wang, J., & Kaufman, R. J. (2007). ATF6alpha optimizes long-term endoplasmic reticulum function to protect cells from chronic stress. Dev Cell, 13(3), 351–364. doi:S1534-5807(07)00266-3 [pii]10.1016/j.devcel.2007.07.005

Xin, Y., Dominguez Gutierrez, G., Okamoto, H., Kim, J., Lee, A. H., Adler, C., & Gromada, J. (2018). Pseudotime Ordering of Single Human beta-Cells Reveals States of Insulin Production and Unfolded Protein Response. Diabetes, 67(9), 1783–1794. doi:10.2337/db18-0365

Yao, C. H., Liu, G. Y., Wang, R., Moon, S. H., Gross, R. W., & Patti, G. J. (2018). Identifying off-target effects of etomoxir reveals that carnitine palmitoyltransferase I is essential for cancer cell proliferation independent of beta-oxidation. PLoS Biol, 16(3), e2003782. doi:10.1371/journal.pbio.2003782

Ying, W. (2008). NAD+/NADH and NADP+/NADPH in cellular functions and cell death: regulation and biological consequences. Antioxid Redox Signal, 10(2), 179–206. doi:10.1089/ars.2007.1672

Yoboue, E. D., Sitia, R., & Simmen, T. (2018). Redox crosstalk at endoplasmic reticulum (ER) membrane contact sites (MCS) uses toxic waste to deliver messages. Cell Death Dis, 9(3), 331. doi:10.1038/s41419-017-0033-4

Zhao, Y., Seefeldt, T., Chen, W., Wang, X., Matthees, D., Hu, Y., & Guan, X. (2009). Effects of glutathione reductase inhibition on cellular thiol redox state and related systems. Arch Biochem Biophys, 485(1), 56–62. doi:S0003-9861(09)00064-2 [pii]10.1016/j.abb.2009.03.001

Zito, E., Chin, K. T., Blais, J., Harding, H. P., & Ron, D. (2010). ERO1-beta, a pancreas-specific disulfide oxidase, promotes insulin biogenesis and glucose homeostasis. J Cell Biol, 188(6), 821–832. doi:10.1083/jcb.200911086

Zito, E., Melo, E. P., Yang, Y., Wahlander, A., Neubert, T. A., & Ron, D. (2010). Oxidative protein folding by an endoplasmic reticulum-localized peroxiredoxin. Mol Cell, 40(5), 787–797. doi:10.1016/j.molcel.2010.11.010

